# GWAS of brain volume on 54,407 individuals and cross-trait analysis with intelligence identifies shared genomic loci and genes

**DOI:** 10.1101/613489

**Authors:** Philip R Jansen, Mats Nagel, Kyoko Watanabe, Yongbin Wei, Jeanne E Savage, Christiaan A de Leeuw, Martijn P van den Heuvel, Sophie van der Sluis, Danielle Posthuma

**Affiliations:** Department of Complex Trait Genetics, Center for Neurogenomics and Cognitive Research, Amsterdam Neuroscience, Vrije Universiteit, Amsterdam, The Netherlands; Department of Clinical Genetics, Section Complex Trait Genetics, Amsterdam Neuroscience, Vrije Universiteit Medical Center, Amsterdam UMC, Amsterdam, the Netherlands; Department of Child and Adolescent Psychiatry, Erasmus University Medical Center, Rotterdam, the Netherlands; Connectome Group, Department of Complex Trait Genetics, Center for Neurogenomics and Cognitive Research, Amsterdam Neuroscience, Vrije Universiteit, Amsterdam, the Netherlands

**Author notes:** These authors contributed equally to this work (Philip R Jansen, Mats Nagel). These authors jointly supervised this work (Sophie van der Sluis, Danielle Posthuma). Correspondence should be addressed to: Danielle Posthuma: Department of Complex Trait Genetics, Vrije Universiteit Amsterdam, De Boelelaan 1085, 1081 HV, Amsterdam, The Netherlands.

## Abstract

The phenotypic correlation between human intelligence and brain volume (BV) is considerable (*r*≈0.40), and has been shown to be due to shared genetic factors^1^. To further examine specific genetic factors driving this correlation, we present genomic analyses of the genetic overlap between intelligence and BV using genome-wide association study (GWAS) results. First, we conducted the largest BV GWAS meta-analysis to date (N=54,407 individuals), followed by functional annotation and gene-mapping. We identified 35 genomic loci (27 novel), implicating 362 genes (346 novel) and 23 biological pathways for BV. Second, we used an existing GWAS for intelligence (N=269,867 individuals^2^), and estimated the genetic correlation (*r_g_*) between BV and intelligence to be 0.23. We show that the *r_g_* is driven by physical overlap of GWAS hits in 5 genomic loci. We identified 67 shared genes between BV and intelligence, which are mainly involved in important signaling pathways regulating cell growth. Out of these 67 we prioritized 32 that are most likely to have functional impact. These results provide new information on the genetics of BV and provide biological insight into BV’s shared genetic etiology with intelligence.

## GWAS of brain volume

To identify SNPs associated with brain volume (BV), we first performed a GWAS (**Supplementary Fig. 1**) using data of 17,062 participants from the UK Biobank^3^, with BV estimated from structural (T_1_-weighted) magnetic resonance imaging (MRI) by summing total gray and white matter volume, and ventricular cerebrospinal fluid volume (**Supplementary Fig. 2**). These results were then meta-analyzed with GWAS results from two previously published studies^4,5^: one on intracranial volume (ICV) from the ENIGMA-CHARGE^4^ collaboration (N=21,875 individuals), and a second one on head circumference, a proven proxy of BV^6–8^, from the Early Growth Genetics (EGG) consortium^5^ (N=10,768 individuals). This led to a total sample size of 54,407 unrelated Europeans **(Supplementary Fig. 3; Supplementary Methods 1.1-1.2; Supplementary Table 1**). Linkage disequilibrium score regression (LDSC^9^; **Online Methods**) showed high concordance of SNP associations between the three samples (UKB and ENIGMA-CHARGE: *r_g_*= 1.04, SE=0.07; UKB and EGG: *r_g_*=0.80, SE=0.14; ENIGMA-CHARGE & EGG: *r_g_*=0.71 SE=0.15; **Supplementary Table 2**), justifying subsequent meta-analysis. Sample-size weighted fixed-effects meta-analysis was carried out using METAL^10^ (**Online Methods**) resulting in 46 linkage disequilibrium (LD) independent lead SNPs (*r*^2^<0.1), residing in 35 genomic loci (**Fig. 1a; Supplementary Table 3; Supplementary Fig. 4**), representing 4,583 genome-wide significant SNPs (**Fig. 1b, Supplementary Table 4**) associated with our measure of BV. Of these 35 loci, 27 were novel compared to the latest GWAS study of intracranial volume^4^. To see if our results were driven by only one of the three samples or were supported by all three, we examined the direction of effect of all genome-wide significant (GWS) SNPs across the three individual cohorts. We found high concordance: 99.5% of the 4,583 SNPs had the same direction of effect in all three cohorts. Of the 4,583 SNPs that were GWS in the BV meta-analysis, 15% was GWS in at least two of the individual cohorts, while 77% (19.7%) had a *P*-value <0.05 in at least two (all three) cohorts. The SNP-heritability (*h*^2^_*SNP*_) of BV estimated by LDSC was 0.24 (SE=0.02), and the LDSC intercept approximated 1 (1.037, SE=0.009), suggesting that the inflation in test statistics (Lambda λ_GC_=1.23) was largely due to polygenicity and not to unaccounted for population stratification^11^. Functional annotation of 5,802 ‘candidate’ SNPs (i.e., SNPs in the risk loci with a GWAS *P*-value of *P*<10^−5^ and LD *r*^2^ >0.6 with one of the independent significant SNPs; **Online Methods**) carried out in FUMA^12^ showed that these SNPs were most abundant in intronic (n=2,879, 49.6%) or intergenic regions (n=l,131, 19.5%), these proportions were significantly enriched (compared to 37.5%, *P*=1.63×10^−81^) or depleted (compared to 45.6%, *P*<1×10^−100^) relative to all SNPs included in the meta-analysis, respectively (**Supplementary Fig. 5; Supplementary Table 5**), and 30 (0.5%) were exonic nonsynonymous SNPs (ExNS) altering protein structures of 15 genes (**Supplementary Table 6, Supplementary Results 2.1**). One gene, *SPPL2C*, contained 8 ExNS (all in exon 1, and in the same inversion region). *SPPL2C* codes for the signal peptide peptidase-like 2C, which plays a role in the degradation of signaling peptides in the brain^13^.

**Figure 1.**
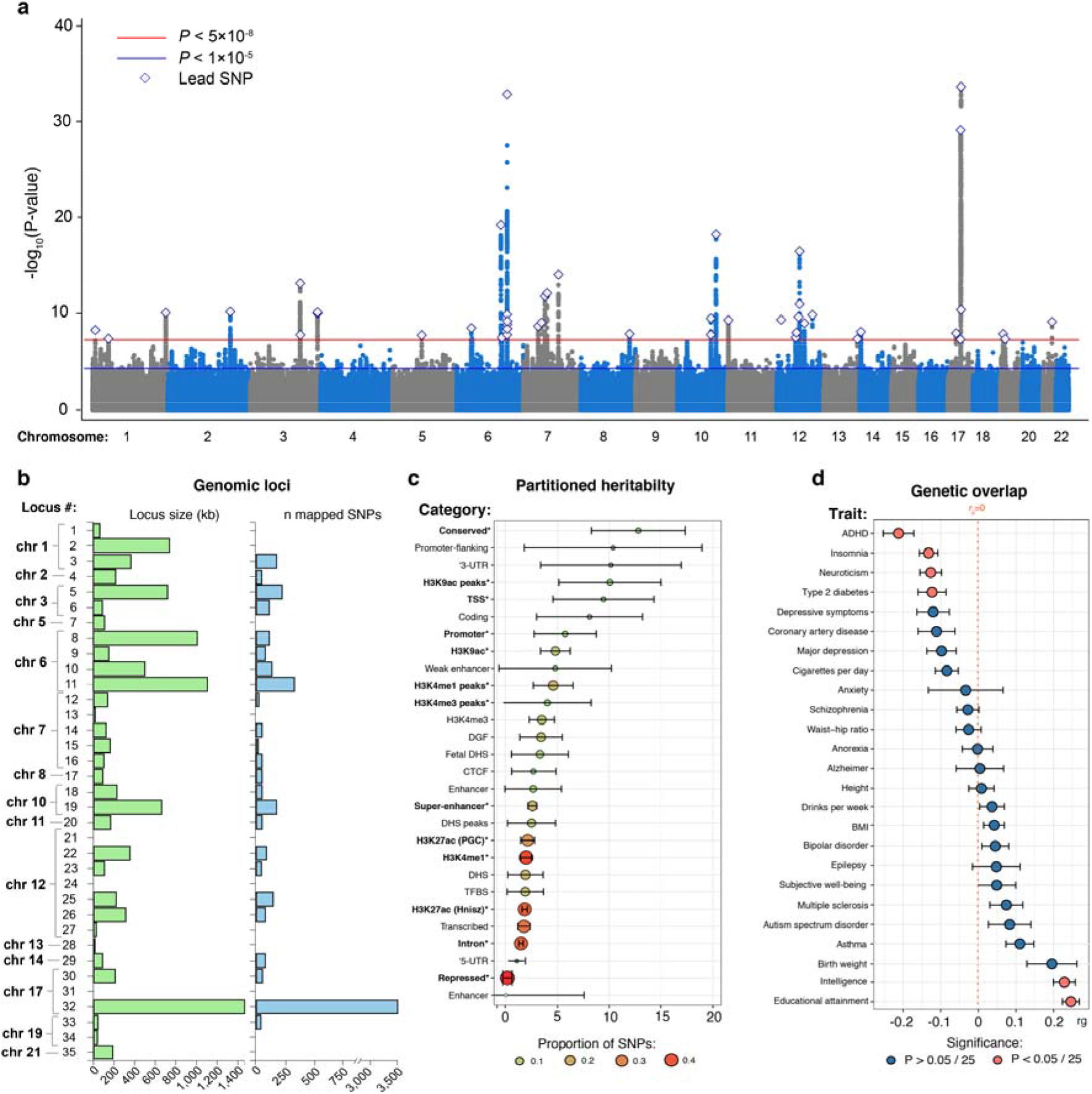
Genome-wide meta-analysis and follow-up analysis of brain volume (BV) in UK Biobank, ENIGMA-CHARGE and Early Growth Genetics consortium. **(a)** Manhattan plot of the GWAS meta-analysis of BV in 54,407 individuals showing the genomic position of each SNP on the *x*-axis and the meta-analysis negative log_10_-transformed *P*-value on the *y*-axis. Independent lead SNPs of each locus are annotated by a diamond. The horizontal red line indicates the genome-wide significance threshold that corresponds to a *P*-value of 5×10^−8^, while the horizontal blue indicates the suggestive threshold of 1×10^−5^. **(b)** Overview of the genomic loci sizes and number of SNPs mapped by each locus. **(c)** Partitioning of the SNP heritability of the BV GWAS meta-analysis by binary SNP annotations using LD Score regression. Enrichment was calculated by dividing the partial heritability of a category by the proportion of SNPs in that category. Significant enrichments, corrected for the number of categories tested (*P*<0.05/28), are highlighted in bold and with an asterisk. **(d)** Genetic correlations with previous traits estimated using LD Score regression. Red dots denote significant genetic correlation after Bonferroni correction (*P*<0.05/25).

To test whether specific functional categories SNP annotations contribute disproportionally to the heritability of BV, we used LDSC to partition the genetic signal over functional SNP categories (**Online Methods**). We observed significant heritability enrichment in 13 SNP categories (**Fig. 1c; Supplementary Table 7**), with the strongest enrichment of SNPs in (evolutionary) conserved regions, (enrichment=12.8, SE=2.3, *P*=1.03×10^−6^, suggesting an 12.8-fold increase in *h*^2^ conveyed by SNPs in these regions), H3K9ac peaks (i.e., specific histone modification that is correlated with active promoters; enrichment=10.1, SE=2.5, *P*=5.58×10^−4^) and transcription start sites (TSS; enrichment=9.5, SE=2.5, *P*=8.48×10^−4^).

## BV gene analyses

To gain insight into which genes may be involved in BV, we mapped the candidate SNPs implicated in the BV meta-analysis to genes, using positional mapping, eQTL mapping, and chromatin interaction mapping as implemented in FUMA^12^, and by performing gene-based association tests as implemented in MAGMA^14^ (**Online Methods, Supplementary Fig. 6**). In total, 362 unique genes were implicated by at least one of these methods (of these, 140 genes were overlapping with one of the 35 identified risk loci). Specifically, the 35 risk loci were mapped to 101 genes based on position, 186 genes by eQTL association, 201 genes through chromatin-chromatin interactions (**Fig. 2a; Supplementary Table 8**), and gene-based association testing identified 70 genes (**Fig. 2b; Supplementary Table 9**). The 362 genes included 17 genes that were replicated from previous studies (5 not replicated), while 345 were novel gene findings (**Supplementary Table 10**). Overall, 16 genes were implicated by all methods (*CDK6, PRMT5, ERBB3, FOXO3, FRZB, FAM49B, PRR13, WBP1L, HAUS4, RBM23, PTEN, RAB5B, MAP3K12, HMGA2, AJUBA, INA*, of which 13 novel gene findings).

**Figure 2.**
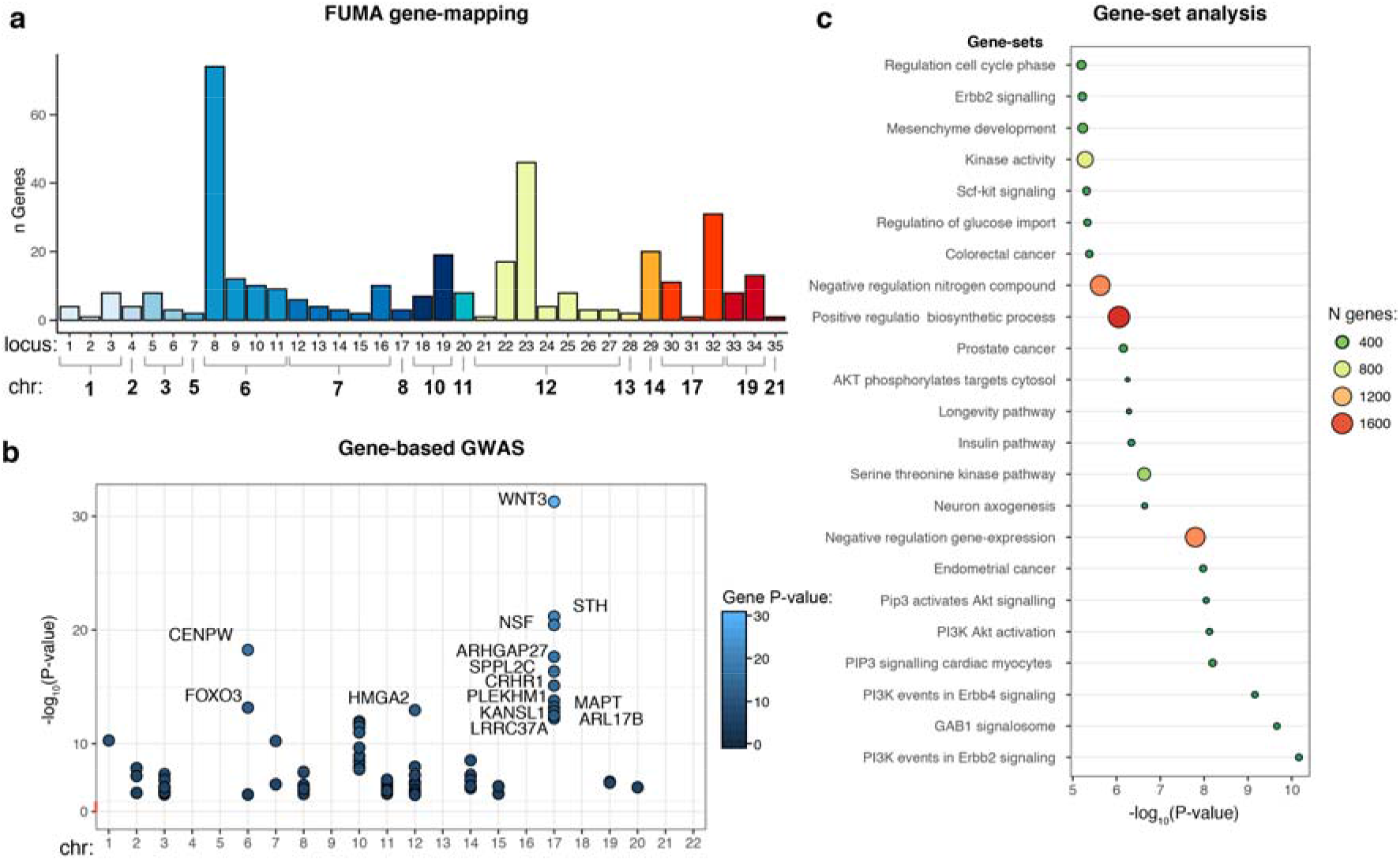
Gene analyses based on the GWAS meta-analysis of BV. **(a)** Number of genes that were mapped by FUMA to each locus. Colors indicate distinct chromosome numbers **(b)** Significant genes in the gene-based association test in MAGMA after Bonferroni correction (*P*<0.05/18,161). The top 15 most significant genes are annotated with the corresponding gene symbols. **(c)** Significant MsigDB gene-sets after Bonferroni correction (*P*<6.36×10^−6^ (=0.05/(7,246+53+565)) showing the negative log_10_-transformed *P*-value on the *x*-axis; size and colors of the circles represent the size of the gene set.

Since monogenic Mendelian disorders are often characterized by abnormal brain development, we performed look-up of the 362 genes implicated in the BV GWAS in the Online Mendelian Inheritance in Man database^15^ (OMIM, **URLs**). We identified 106 monogenic disorders caused by high-penetrance mutations in 88 of the 362 genes (24.3% vs. 17.1% in all genes that were tested in the gene-based analysis, *P*=3.39×10^−4^) implicated in BV, many of which are commonly associated with abnormal brain development (including micro- (*CDK6*) and macrocephalia (*PTEN*)), and abnormal growth (including leprechaunism (*INSR*) and tetra-amelia syndrome (*WNT3*)) (**Supplementary Table 11**).

To identify functional pathways related to BV, we performed gene-set analysis in MAGMA^16^(**Online Methods**). Of the 7,246 tested gene-sets (including canonical pathways and gene ontology gene-sets), 23 were significantly associated after Bonferroni correction (**Fig. 2c; Supplementary Table 12-13**). Among these significant gene-sets were several cell-signaling pathways, including the ErBB2/ErBB4 signaling, the GAB1 signaling pathway and the insulin pathway, and developmental processes such as neuron axogenesis and mesenchyme development. That is, key cell-signaling pathways were implicated that are involved in normal brain development^17^ and involved in several brain developmental abnormalities^18,19^. Pairwise conditional gene-set analysis indicated that several gene-sets (e.g. neuron axogenesis and mesenchyme development) constitute independent associations. (**Supplementary Fig. 7-9, Supplementary Table 14, Supplementary Results 2.4**). Conversely, conditional analyses suggest that several signaling pathways (e.g., PI3K events in ERBB2/ERBB4 signaling, GAB1 signalosome) represent a shared underlying signal, since conditional P-values were substantially higher than the unconditional *P*-values.

Next we aimed to identify tissue categories and neuronal cell-types that are enriched for gene signal of BV, by linking gene *P*-values to gene-expression in 53 tissue-types^20^ and 565 brain cell-types^21^ (**Online Methods**). None of the associations of tissue or cell-types passed our stringent multiple testing correction (i.e. correcting for all tested gene-sets, tissues and cell types, thus 0.05/7,864), but the strongest evidence of association was observed for several types of polydendrocytes, including *TNR-PDGFA-PIK3R3* (*P*=7.46×10^−4^) and *TNR-CSPG5* (*P*=2.53×10^−3^) polydendrocytes (**Supplementary Table 12**). Polydendrocytes are thought to be precursor cells of oligodendrocytes^22^, a cell-type involved in supporting neuronal health and myelinization of the brain^23^.

To determine to what extent our BV GWAS results were the product of a heterogeneous phenotype (structural MRI and head circumference), an additional meta-analysis was performed on a stricter, fully height-adjusted, BV phenotype. The outcomes resembled those of the primary meta-analysis very closely (*r_g_*=0.96, SE=0.007), increasing our confidence in the main results (see **Supplementary Tables 15-22** and **Supplementary Figs. 10-12** for the results of the SNP- and gene-level analysis, which are discussed in **Supplementary Results 2.5**).

## Genetic correlations between BV and other traits

Twin studies have shown substantial genetic overlap between BV and behavioral traits, including intelligence^1,24–26^. We used LDSC to estimate the overlap in genetic signal between BV and 25 brain-related and neuropsychiatric traits for which published summary statistics based on large samples were available (**Online Methods**). Significant genetic correlations were observed between BV and 6 traits: positive genetic correlations with educational attainment (*r_g_*=0.25, SE=0.02, *P*=5.14×10^−28^; **Fig. 1d; Supplementary Table 23**) and intelligence (*r_g_*=0.23, SE=0.03, *P*=2.82×10^−15^), and negative genetic correlations with ADHD (*r_g_*=-0,21 SE=0.04, *P*=1.97×10^−7^), type 2 diabetes (*r_g_*=-0.12, SE=0.04, *P*=9.56×10^−4^), insomnia (*r_g_*=-0,13 SE=0.02, *P*=6.01×10^−8^), and neuroticism (*r_g_*=-0.13, SE=0.03, *P*=1.01×10^−5^), confirming previously reported overlap^4,27, 28^ and establishing a new association with type 2 diabetes.

## Genetic overlap with intelligence

Development of human intelligence has coincided with a strong increase in total size of the brain^29^. Indeed, epidemiological studies have shown overlap in genetic factors between BV and intelligence^1,2^, which we confirmed here: the estimated *r_g_* between BV and intelligence was 0.23 (**Supplementary Tables 2, 23**), closely resembling the estimates of shared genetic factors between intelligence and gray matter volume (*r_g_*=0.29) and white matter volume (*r_g_*=0.24) obtained from twin studies^1^. To gain insight into the specific genetic factors driving this genetic correlation, we explored the overlap in genetic signal between the current BV GWAS meta-analysis and a recent large GWAS meta-analysis of intelligence^2^ (N=269,867).

We first determined whether strongly associated SNPs show the same direction of effect in both traits, by performing a look-up of the lead SNPs of BV in the intelligence GWAS, and vice versa. In line with expectations, we observed a strong sign concordance of the 46 BV lead SNPs in intelligence (sign concordance=89.1%, *P*=3.10×10^−7^, **Fig. 3c**) and weaker but still considerable sign concordance of the 243 lead SNPs of intelligence in BV (sign concordance=62.1%, *P*=1.86×10^−4^, **Fig. 3d**). Similarly, strong deviations from the expected P-value distribution were observed for genes that were significant for BV (70) and intelligence (507) in the gene-based test in MAGMA (**Supplementary Fig. 13**).

**Figure 3.**
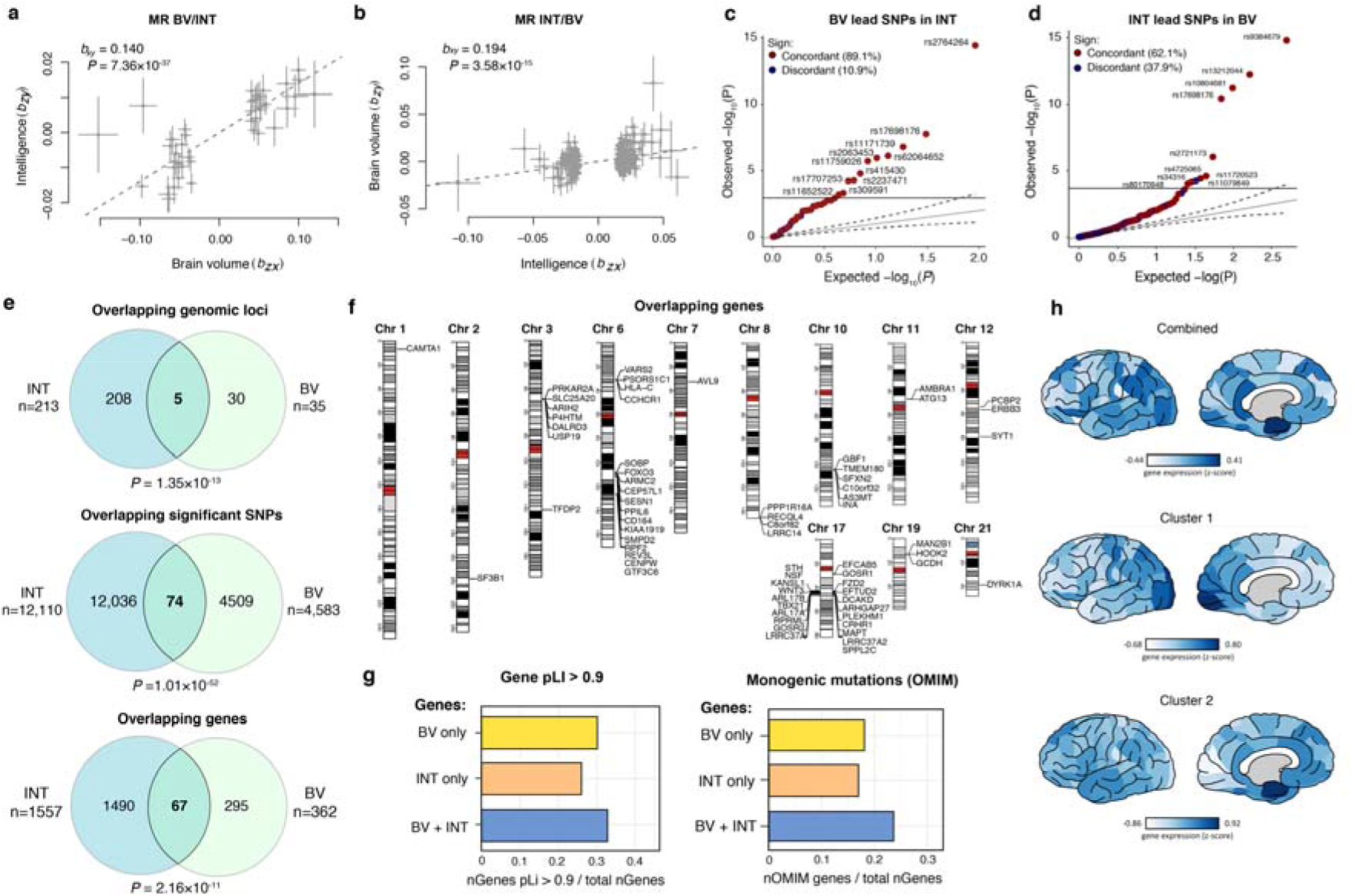
Analyses of the genetic overlap between BV and intelligence. **(a)** Mendelian Randomization analysis, showing the effects and standard errors of each SNP of BV (*b_zy_* on intelligence (*b_zx_*) and the diagonal line showing the estimated causal effect (*b_xy_*) of BV on intelligence. **(b)** Similar to a) but of intelligence on BV. **(c)** Q-Q plot showing the observed and expected P-value distribution of 46 lead SNPs of the BV meta-analysis in intelligence and **(d)** 243 intelligence lead SNPs in BV. The colors denote the concordance (/discordance) of effects between the two traits. **(e)** Venn diagrams showing the number of overlapping loci, significant SNPs and genes implicated in BV and intelligence by FUMA and MAGMA. **(f)** Karyogram showing the genomic position of genes that were implicated in the GWAS of both BV and intelligence. **(g)** Comparison of the pLI score (representing tolerance to loss-of-function variants) and fraction of genes involved in developmental disorders between BV-only genes, intelligence-only genes (INT), and genes associated to bot traits (BV+INT). **(h)** Cortical gene expression patterns (colors indicate rank scores) across 57 cortical areas of the 67 genes that overlap between BV and intelligence, and the two clusters.

To explore whether BV and intelligence are causally related, we carried out GWAS summary statistics-based Mendelian Randomization analyses, using the Generalized Summary-data-based Mendelian Randomization package^30^ (GSMR; **Online Methods**), essentially using independent lead SNPs as instrumental variables (LD: *r*^2^<0.1). GSMR analyses demonstrated a directional effect of BV on intelligence (*b_xy_*=0.140, SE=0.011, P=7.36×10^−37^; **Fig. 3a**) and a less strong yet still highly significant directional effect of intelligence on BV (*b_xy_*=0.194, SE=0.025, P=3.58×10^−15^; **Fig. 3b**), suggesting a bidirectional association between these phenotypes, in line with previous reports^2^. We do note, however, that GSMR makes several strong assumptions, such as the absence of a third, mediating, factor, and no horizontal or vertical pleiotropy^31^, which may not always hold, and that the current results should be interpreted conditional on these assumptions.

To investigate whether specific SNPs or genes could be identified that drive the genetic overlap between intelligence and BV, we performed several cross-trait analyses of SNPs and genes significantly implicated in both traits. We observed physical overlap in five out of the 35 genomic loci for BV, and overlap in genes implicated by FUMA (n_genes_=45) and MAGMA (n_genes_=23), resulting in 67 unique overlapping genes (**Fig. 3e-f; Supplementary Table 24**). Conversely, we identified 295 genes for BV that were not significantly associated to intelligence (**Fig. 3e**). Lookup of gene functions of the 67 overlapping genes in the online GeneCards^32^ repository (**URLs**) showed strong involvement of these genes in a wide variety of cellular processes and key factors in cell division (**Supplementary Results 2.6**). When comparing the probability of being loss-of-function (LoF) intolerant (pLI score >0.9) between overlap genes and genes observed for only one of these traits, we observe a slightly higher fraction of genes being LoF intolerant in the genes that play a role in both traits (32.8% (shared) vs. 26.0% (intelligence) and 30.2% (BV), **Fig. 3g**,). The mutation intolerance of these overlapping genes is further demonstrated by a high fraction of genes in this category that are associated with monogenic disease (23.8%, **Fig. 3g**) in the OMIM database), compared to BV-only (18.9%) and intelligence-only (17.2%) genes. The finding that the overlapping genes, that are involved in multiple key cellular processes, are overall less tolerant to (LoF) mutations, implies that their unperturbed functioning is crucial to a well-functioning human brain.

In order to determine whether the set of genes that overlap between BV and intelligence are randomly expressed across cortical areas or show similar expression profiles across cortical brain areas, we examined their cortical gene expression profiles using data obtained from the Allen Human Brain Atlas^33^ (AHBA). We compared the gene expression profile of the 67 genes related to both BV and intelligence to that of genes associated to either or both of the traits (n_genes_=1,852; **Fig. 3e**). Specifically, using permutation analysis we compared the expression of these 67 genes across 57 brain regions to that of 10,000 randomly selected, equally large, sets of genes drawn from the 1,852 genes related to either or both of the traits. Although we observed overexpression of the 67 overlapping genes in the anterior part of the fusiform (*P*=8.12×10^−3^) and the posterior part of the inferior temporal cortex (*P*=7.82×10^−3^), none of the associations survived a stringent Bonferroni correction (*P*<8.77×10^−4^ (=0.05/57); **Supplementary Table 25**). To examine whether clusters of more homogeneously expressed genes exist within the 67 overlapping genes, we additionally performed clustering analysis on the correlations between the expression profiles of genes, aiming to maximize intra-cluster cohesion, while at the same time maximizing differences between clusters (see **Online Methods**). Two clusters were identified (**Supplementary Results, Supplementary Table 25**) but their expression profiles were not significantly different from those of random sets. Interaction analyses in MAGMA (see **Online Methods**), testing whether gene expression in specific brain regions contributed significantly to the genetic relationship between BV and intelligence, also turned out negative (**Supplementary Table 26**). Based on these results, we conclude that the set of overlapping genes do not seem to be expressed in any particular cortical region.

To prioritize those genes from the 67 overlapping genes, that are more likely to have causal effects on both BV and intelligence, we filtered the set of common genes based on either one of three conditions: 1) the gene contained an ExNS SNP, 2) the gene was part of one of the significant gene sets for BV or intelligence, and 3) the GWAS signal of either trait colocalized with eQTL signals, using COLOC^34^ (colocalization of GWAS and eQTL signals is compatible with the hypothesis of a common causal variant; see **URLs; Supplementary Table 24**). After filtering the 67 genes that were associated with both BV and intelligence, a selection of 32 potentially functionally interesting genes (17 of which did not physically overlap with each other) remained. For example, we prioritized two genes on chromosome 3 (out of 6 associated genes in the same region) that may proof particularly interesting candidates for functional follow-up aimed at characterizing the genetic relation between BV and intelligence: *USP19* was selected because it contains an ExNS variant that was significantly associated to intelligence, and *TFDP2* because it was part of a significantly associated gene set for BV.

Interestingly, conditional gene-set analyses (where we conditioned gene-based signal in BV on the gene-based Z-scores for intelligence, and vice versa) indicated that the gene sets identified for BV (23) and intelligence (6) are trait specific, since the association remains virtually unchanged when conditioning on the other trait (**Supplementary Table 27**). Fisher’s exact tests showed that the 67 overlapping genes were not significantly enriched in the 29 gene sets associated to BV or intelligence (**Supplementary Table 28**). However, several genes related to intelligence were located in gene-sets observed for intelligence, including *TFDP2, CDKN1A, FOXO3 and VARS1* (see **Fig. 4**), suggesting that, although no single gene-set was shared between BV and intelligence, several genes associated with intelligence are located along important signaling pathways implicated in BV.

**Fig 4.**
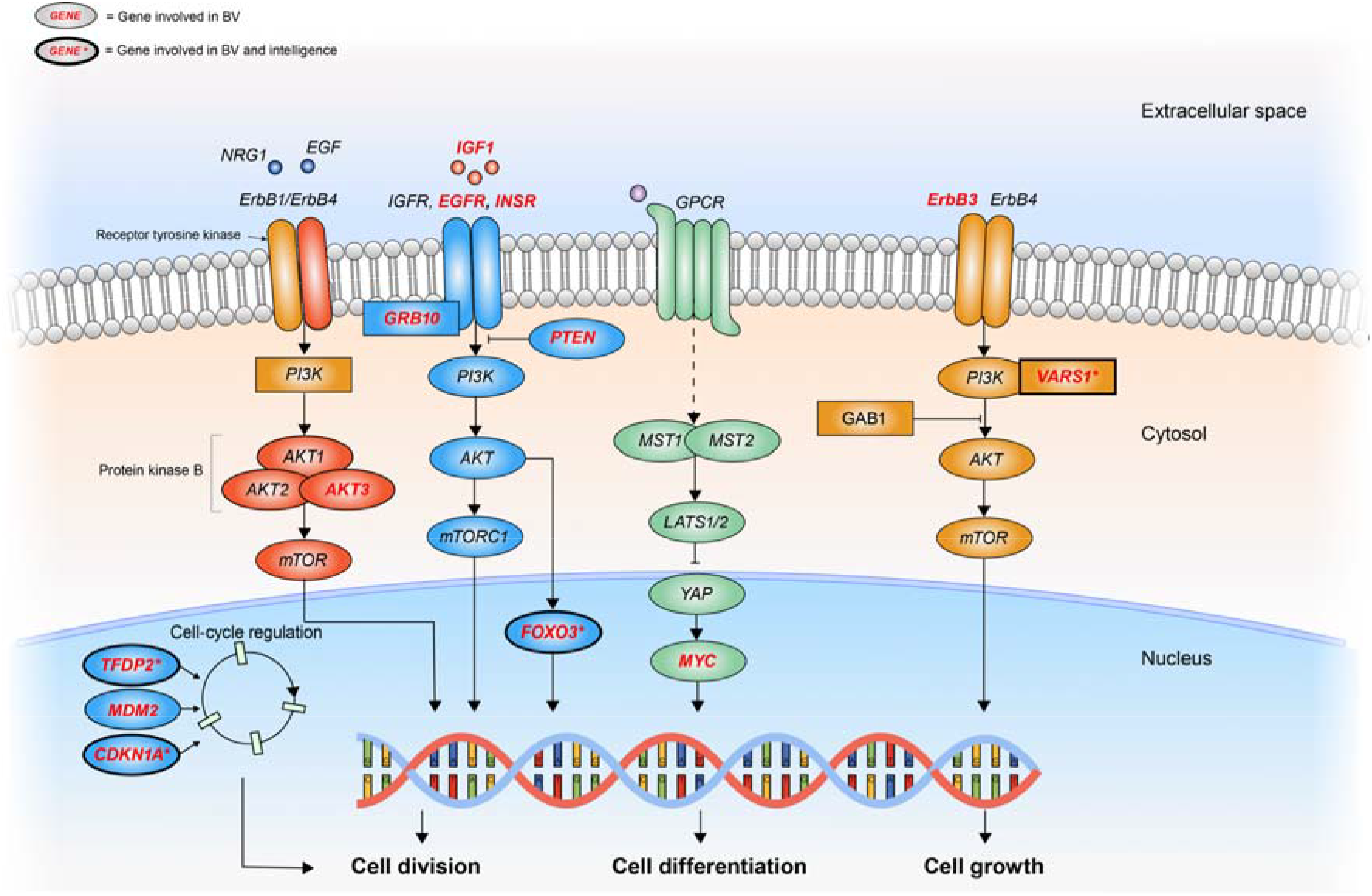
Schematic representation of cellular signaling pathways involved in BV. Overview of cellular signaling pathways and implicated genes within these pathways that were mapped by the GWAS meta-analysis of brain volume (BV) or overlapped with the genes that were mapped by the GWAS of intelligence. Genes in highlighted in red were involved in the meta-analysis of BV, whereas genes highlighted in red in a thick-outlined oval denoted by an asterisk were observed in both the BV and intelligence GWAS.

In conclusion, we studied the genetics of BV and report 29 novel loci and 346 novel genes for BV. In exploring the genetic overlap between BV and intelligence, we identified 67 genes that are associated to both traits, i.e., contributing to the genetic overlap between these traits. Of the 67 overlapping genes, we prioritized 32 genes that might be best candidates for functional follow-up aimed at gaining biological insight into the overlap between BV and intelligence. We did not identify specific biological characteristics or mechanisms that differentiate the 67 overlapping genes from the full set of genes that was investigated, except for an enriched intolerance to (LoF) mutations. We expect that increasing the sample size, and thus the statistical power, for brain volume GWAS will help in further dissecting the shared nature of brain volume and intelligence. These findings are a first step towards understanding of the genetic architecture of BV and its shared etiology with intelligence.

## ONLINE METHODS

### Samples & phenotypes

#### UK Biobank - Total brain volume

The UK Biobank (UKB) constitutes a large data set, combining a wide range of phenotypes with genetic and imaging information. Here, we used processed data of a subset of N=21,407 individuals who underwent a magnetic resonance imaging (MRI) procedure: data were released in the third quarter of 2018. After filtering on quality of the imaging results, relatedness, European ancestry and the availability of relevant covariates (discussed in more detail below), we arrived at a final sample size of N=17,062. The total brain volume (TBV) phenotype was approximated as follows: TBV = white matter volume + grey matter volume + cerebrospinal fluid volume. The UKB obtained ethical approval from the National Research Ethics Service Committee North West–Haydock (reference 11/NW/0382), and all study procedures were performed in accordance with the World Medical Association for medical research. The current study was conducted under UKB application number 16406.

#### ENIGMA-CHARGE collaboration - Intracranial volume

We used GWAS summary statistics on intracranial volume obtained in a collaboration between the Cohorts for Heart and Aging Research in Genomic Epidemiology (CHARGE) and Enhancing NeuroImaging Genetics through Meta-Analysis (ENIGMA)^35^. ICV was calculated from brain MRI data collected in 46 individual cohorts. Given the strong correlation between brain volume and height (r=0.55 in UK Biobank) we considered height an important covariate to include in GWAS on volumetric brain measures. Therefore, we used mtCOJO to correct for height, by conditioning on height using UKB-derived sumstats (this procedure is described in more detail below). Aside from the full summary statistics (N=26,577), we also obtained summary statistics for a subset of individuals for who ‘height’ was available as a covariate (N=21,875). Details on the individual cohorts included in the ENIGMA-CHARGE collaboration are described elsewhere^35^. All participants were of European descent and provided informed consent. All studies included in the ENIGMA-CHARGE collaboration were approved by their local institutional review board or local ethics committee. Details on genotyping, imputation and analysis procedures can be found in the original publication^35^.

#### EGG consortium

In contrast to the two samples discussed above, the GWAS summary statistics obtained from the Early Growth Genetics (EGG) consortium are based on genetic analysis of an indirect measure of brain volume, i.e., head circumference in infancy. These summary statistics were generated by meta-analyzing seven population-based European studies, resulting in a total sample size of N=10,678. All studies included in the meta-analysis were approved by the local ethics committees, and informed (or parental) consent was obtained from all participants. Data collection, genotyping, imputation and analysis procedures were described in detail elsewhere^36,37^. Since not all individual studies that were included in the EGG consortium corrected for height, we first conditioned the summary statistics on height using a procedure similar to that described for the ENIGMA-CHARGE data.

### Genotyping, imputation & quality control

#### UK Biobank

The genotype data that we used was released by the UKB in March 2018, and concerns an updated version of data released earlier (July 2017). Details on the collection and processing of the genotype data are described elsewhere^3^. To summarize, the UKB genotyped in total 489,212 individuals on two custom made SNP arrays (UK BiLEVE Axiom^™^ array covering 807,411 markers (n=49,950) and UK Biobank^™^ Axiom array covering 825,927 markers (n=438,427), both by Affymetrix). The genotype arrays shared 95% of marker content. Quality control executed by the UKB team resulted in a total of 488,377 individuals and 805,426 unique markers in the released data. In the version of the data used for the current study (release March 2018), genotypes were phased and imputed by the UKB team to a combined reference panel of the Haplotype Reference Consortium and the UK10K. Finally, imputed and quality-controlled genotype data was available for 487,422 individuals and 92,693,895 genetic variants. Prior to analysis, imputed variants were converted to hard call using a certainty threshold of 0.9. In our own quality control procedure, we excluded SNPs with a low imputation score (INFO score <0.9), low minor allele frequency (MAF<0.005) and high missingness (>0.05). Furthermore, indels positioned in the same chromosomal location were excluded. This resulted in a total of 9,203,453 SNPs used for downstream analysis.

To prevent population stratification from biasing the results, we only included individuals from European descent in our analyses. To this end, we projected principal components (PC’s) from the 1000 Genomes reference populations^38^ onto the called genotypes available in the UKB data. Participants for whom the projected principal component score was closest (using the Mahalanobis distance) to the average score of the European 1000Genomes sample^39^ were considered to be of European descent. Participants having a Mahalanobis distance >6 S.D. were excluded from further analysis. In an additional quality control step, we excluded participants that 1) had withdrawn their consent, 2) were related according to the UKB team (i.e., subjects with most inferred relatives, 3^rd^ degree or closer, were removed until no related subjects were present), 3) reported a gender that did not match their genetic gender, or 4) showed sex-chromosome aneuploidy. After filtering availability of imaging data, and MRI scan quality, 17,062 individuals remained for analysis.

### GWAS of total brain volume in UKB

The genome-wide association analyses (GWAS) of total brain volume (BV) in the UKB data was conducted in PLINK^40,41^, using a linear regression model with additive allelic effects. In order to correct for potential subtle population stratification effects, we included 10 genetic PC’s as covariates. Genetic PC’s were computed using FlashPC2^42^ in the QC’ed subset of unrelated European subjects, retaining only independent (*r*^2^<0.1), relatively common (MAF>0.01) and genotyped or very high imputation quality (INFO=1) SNPs (n=145,432 markers). Additional covariates included in the analysis were: age, sex, genotype array, assessment center, standing height, and the Townsend deprivation index.

### Conditional GWAS (mtCOJO)

We used multi-trait-based conditional & joint analysis using GWAS summary data (mtCOJO^30^; see **URLs**) to conduct conditional GWAS analyses of our traits of interest. The mtCOJO method is integrated in the GCTA^43^ software, and requires summary-level GWAS data to perform a GWAS analysis of phenotype A, conditioned on phenotype B. In the current study, we performed two conditional GWASs, aiming to correct for height in two samples measuring ICV (ENIGMA-CHARGE) and head circumference (EGG), respectively. First, we ran a conditional GWAS for ICV in the full ENIGMA-CHARGE sample, conditioning on UKB-derived summary statistics for standing height. Second, we ran a similar conditional analysis for head circumference in the EGG sample, again conditioning on UKB-derived summary statistics for standing height. Results from the conditional GWASs were subsequently included in the main GWAS meta-analysis of brain volume described in the following section.

### GWAS meta-analysis of brain volume

Before carrying out the meta-analysis of brain volume (BV), we performed additional filtering and prepared the summary statistics for each of the individual cohorts. First of all, within the ENIGMA-CHARGE data, SNPs that were available for *N*<5,000 were excluded. We then performed similar filtering on the EGG data. Secondly, in the UKB data we identified some instances where there were multiple SNPs on the same position, whilst having different alleles. These SNPs were excluded from further analysis. Lastly, we aligned the allele coding of indels in the ENIGMA-CHARGE data to match the coding in the UKB data. A number of indels that was not present in the UKB data were excluded from analysis, since we used the UKB data as reference data for downstream analyses. The summary statistics from the UKB, ENIGMA-CHARGE and the EGG consortium showed strong genetic correlations (*r_g_* ranging between 0.71 and 1; see **Supplementary Table 2**), supporting our choice for a meta-analytic approach.

Using a sample-size weighted *z*-score method in METAL^10^ we combined the GWAS on total brain volume in the UKB data (N=17,062), the GWAS on intracranial volume (conditioned on height) in the ENIGMA-CHARGE data (N=26,577), and the GWAS on head circumference (conditioned on height) in the EGG data, resulting in a total sample size of N=54,407 (see **Supplementary Fig. 1**).

To validate our results, we conducted a second meta-analysis, using a stricter phenotype, and only summary statistics from GWASs in which height was directly included as a covariate (rather than conditioning on height using mtCOJO afterwards). In this stricter meta-analysis, we thus combined the full summary statistics from the UKB (N=17,062) with a subset of the ENIGMA-CHARGE collaboration for which information on height had been available and included as a covariate in the GWAS (N=21,875), resulting in a total sample size of N=38,937. The results of this validation meta-analysis are thus based on a purer BV phenotype (i.e., directly corrected for height and excluding head circumference as a proxy for BV). However, given the high genetic correlations between the two meta-analyses for BV (*r_g_*=.957, SE=0.007) and the larger sample size, we expect statistical power to be greater in the primary meta-analysis including all data. This larger meta-analysis was thus used for all subsequent analyses.

### Intelligence - GWAS summary statistics

We used recently published GWAS meta-analysis summary statistics of intelligence^2^ to study the genetic overlap with brain volume. Data collection procedures and methods are described in detail elsewhere^2^. In comparing the results of BV and intelligence, we updated some of the downstream analyses for intelligence (e.g., we used FUMA version 1.3.2 instead of 1.3.0, we explored a slightly different collection of gene sets, and used GTEx data v7 instead of v6.1). Therefore, gene-based findings presented in the current study may deviate slightly from those presented in the original study^2^. We applied Bonferroni correction for multiple testing to the meta-analytic SNP *P*-values to identify to intelligence associated genetic variants.

### Genomic risk loci and functional annotation

#### SNP annotation

We used FUMA^12^ (v1.3.2; see **URLs**) for the functional annotation of the BV meta-analysis results. FUMA is an online platform that takes GWAS summary statistics as input, and subsequently annotates, prioritizes, and visualizes the results.

Prior to defining genomic risk loci, FUMA identifies variants that are genome-wide significant (5×10^−8^), and independent (*r*^2^<0.6) as *independent significant SNPs*. In the next step, *independent significant SNPs* that are independent from each other at *r*^2^<0.1 are denoted lead SNPs. Genotypes from the UKB were used as reference data to infer LD. Finally, FUMA characterizes genomic risk loci by merging LD blocks that are located close to each other (<250 kb apart). Thus, it is possible that one genomic risk locus contains multiple *independent significant SNPs* or lead SNPs.

In order to obtain information on the functional consequences of SNPs on genes, FUMA performs ANNOVAR^44^ gene-based annotation using Ensembl genes (build 85) for all SNPs in LD (*r*^2^>0.6) with one of the *independent significant SNPs* and having an association *P*-value lower 1×10^−5^. Additionally, CADD scores^45^, Regulome DB scores^46^, and 15-core chromatin state^47,48^ are annotated to SNPs by matching chromosome position, reference, and alternative alleles. CADD scores can be used to prioritize genetic variants that are likely to be pathogenic and/or deleterious (CADD scores >12.37 suggest a SNP is deleterious). The score is a single measure combining various annotations, and has been shown to correlate with pathogenicity, disease severity, and experimentally measured regulatory effects and complex trait associations^49^. RegulomeDB scores^46^ characterize SNPs by their likelihood to have a regulatory functions (with lower scores indicating higher probability of regulatory function). Scores range from 7, meaning that there is no evidence of the variant having a regulatory function, to 1a, meaning that a variant is likely to affect binding and is linked to expression of a gene target^46^. Chromatin state was predicted by ChromHMM^48^ for 127 cell types, using 15 states to classify and describe variants.

#### Gene mapping

Using FUMA^12^, all SNPs in genomic risk loci that were genome-wide significant (*P*<5×10^−8^) or were in LD (*r*^2^>0.6) with one of the *independent significant SNPs*, were mapped to genes. SNPs could be annotated to a gene by either of three strategies. First, positional mapping maps SNPs to protein coding genes based on physical proximity (i.e., within 10 kb window). Second, eQTL mapping maps SNPs to genes whose expression is associated with allelic variation at the SNP level. Information on eQTLs was derived from 3 publicly available data repositories; GTEx^50^ (v7), the Blood eQTL browser^51^ and the BIOS QTL browser^52^. This strategy maps SNPs to genes up to 1 Mb apart (*cis*-eQTLs). We applied a false discovery rate (FDR) of ≤0.05 to limit the results to significant SNP-gene pairs. Third, SNPs were mapped to genes based on significant chromatin interactions between promoter regions of genes (250 bp up- and 500 bp downstream of the transcription start site (TSS)) and a genomic region in a risk locus. In contrast to eQTL mapping, and in the absence of a distance boundary, chromatin interaction mapping may involve long-range interactions. The resolution of chromatin interactions was defined as 40 kb, and hence, interaction regions may comprise multiple genes. In order to prioritize genes implicated by chromatin interaction mapping, information on predicted enhancers and promoters in 111 tissue/cell types from the Roadmap Epigenomics Project^53^ was integrated. We used FUMA to filter on chromatin interactions for which one interaction region overlapped with predicted enhancers, and the other with predicted promoters 250 bp up- and 500 bp downstream of the TSS site of a gene. At the time of writing, FUMA contained Hi-C data of 14 tissue types from the study of Schmitt et al. (2016)^54^. An FDR of 1×10^−5^ was used to define significant interactions.

### Gene-based analysis

Genome-wide gene-based analysis (GWGAS) has the potential to identify genes associated to a trait of interest despite the genetic signal of individual SNPs in or nearby the gene not reaching genome-wide significance in SNP-based analyses. In contrast, a gene harboring a few strongly associated SNP and many SNPs that show only very weak association may not be implicated by gene-based analysis. So in addition to gene-mapping in FUMA, we conducted gene-based analysis in MAGMA^16^ to assess the joint effect of all SNPs within all 19,427 protein-coding genes included in the NCBI 37.3. MAGMA requires as input the *P*-values derived from SNP-based analyses, in this case the BV meta-analysis results. All SNPs in our BV meta-analysis were annotated to genes, resulting in 18,161 and 18,162 genes that contained at least one SNP in the BV meta-analysis and the validation meta-analysis, respectively. Besides SNPs located within a gene, we also included SNPs lying within 2 kb before and 1 kb after the TSS of the gene. We used Entrez ID as the primary gene ID. MAGMA’s gene-based analysis uses a multiple linear principal component regression, where an *F*-test is used to compute the gene *P*-value. The model takes linkage disequilibrium between SNPs into account. Genes were considered to be genome-wide significantly associated if the *P*-value survived a Bonferroni correction for multiple testing (0.05/number of genes tested: *P* < 2.75×10^−6^).

### Gene-set and tissue expression analysis

#### Functional gene-sets

Gene-set analysis was performed using MAGMA^16^, testing 7,246 predefined gene sets in an exploratory fashion. Selected gene sets included canonical pathways (n=1,329) and gene ontology (GO) gene sets (*n*=5,917). All gene sets were obtained from the Molecular Signatures Database (MSigDB, version 6.0; see **URLs**). For all gene-set analyses, competitive, rather than self-contained, P-values are reported. Competitive gene-set analysis tests whether the joint association of genes in a gene set with the phenotype of interest is stronger than that of a randomly selected set of genes of the same size. This approach provides stronger evidence for association of the gene set compared to a self-contained test, where the joint association of genes in a gene set with the phenotype is tested against the null hypothesis of no effect.

#### Gene-expression analysis

To assess whether genes associated to our traits of interest are disproportionately expressed in certain tissue- and cell-types, we applied MAGMA’s gene-expression analysis to investigate associations with several gene expression profiles. First, we tested tissue gene-expression in 53 different tissue types obtained from the GTeX portal (v.7; see **URLs**), which include gene-expression data from 13 brain tissue types^20^. Secondly, we tested gene-expression in 565 distinct adult mouse brain cell-types from Dropviz^21^. These data were collected through the use of the Drop-seq technique^55^ by assessing RNA expression in 690,000 individual cells from 9 brain regions of the adult mice brain, which were subsequently grouped to 565 transcriptionally distinct groups of cell-types.

Gene-set and gene-expression analyses were Bonferroni corrected for the total number of genesets, tissue types and single-cell types tested in MAGMA (*P* < 6.36×10^−6^ (=0.05/(7,246+53+565)).

#### Conditional gene-set analyses

In order to gain more insight in the genetic pathways associated to BV and intelligence, we performed conditional gene-set analyses using MAGMA^16^. Conditional analyses were conducted with the aim of identifying MsigDB gene sets that represent independent associations (i.e., in a regression-based framework, we assessed the association between our trait of interest (e.g., BV) and a gene set, conditional on another trait (e.g., intelligence)). Specifically, we determined which gene set associations remain for BV when we condition on intelligence, and vice versa. This approach provides information on whether gene sets are uniquely associated to BV or intelligence, or rather, shared between both traits. For example, if the *P*-value of association between a gene set and BV increases when conditioning on intelligence (i.e., less significant), then this suggests that the gene-set association is likely shared between both traits, while if the *P*-value of a gene-set and BV is unaffected by conditioning on intelligence, then this implies that the association is specific to BV, i.e., not shared with intelligence. In addition, we conducted pairwise conditional gene-set analysis for all 23 gene-sets that were significantly associated to BV.

### SNP-based heritability and genetic correlations

We used linkage disequilibrium score regression (LDSC)^11^ to estimate the proportion of phenotypic variance that can be explained by common SNPs, a statistic known as SNP-based heritability, *h^2^_SNP_*. We used precomputed LD scores that were calculated using 1000 Genomes European data (see **URLs**).

Genetic correlations (*r_g_*) between the signal from our BV meta-analysis, intelligence^2^, and 24 psychiatric, behavioral and lifestyle-related traits for which summary-level data were available, were also calculated using LDSC^9,11^. Genetic correlations for which the *P*-value survived the correction for multiple testing (Bonferroni-corrected *P* < 0.002 (=0.05/25)) were considered significant.

### Partitioned heritability

In order to determine whether some functional categories of the genome contribute more than others to the SNP-heritability (*h^2^_SNP_*) of BV, we performed stratified LDSC^56^. Using this method, we calculated whether any of the 28 specific genomic categories (see **URLs**) included in the analysis was enriched for SNPs that contribute to *h^2^_SNP_*. Enrichment here is defined as the proportion of *h^2^_SNP_* in a given category divided by the proportion of SNPs in that category (e.g., if enrichment in intronic regions is 4.75, this indicates that this functional category is responsible for a 4.75-fold higher contribution to *h^2^_SNP_* compared to all tested SNPs).

### Mendelian randomization analysis

Mendelian randomization (MR) analysis was performed using Generalized summary-data-based Mendelian randomization (GSMR^30^; see **URLs**). The main goal was to examine whether the genetic correlation between brain volume and intelligence (*r_g_*=0.23) might be explained by directional effects. Analyses were conducted using forward and reverse GSMR, testing for uni- and bidirectional effects between all BV and intelligence.

### Linking expression of overlapping genes for brain volume and intelligence to specific brain regions

To identify brain regions that are associated with genes that are observed in BV and intelligence, we first performed gene-mapping with MAGMA and FUMA based on the summary statistics of the GWAS of each trait separately, after which we extracted the set of genes implicated by any of the gene-mapping strategies in both traits.

Subsequently, we extracted the cortical gene-expression profile for each of the overlapping genes from the Allen Human Brain Atlas^33^ (AHBA), which describes gene-expression data per gene across distinct cortical areas. An expression profile of the set was obtained by taking the average expression across the 67 genes. We also performed clustering on the individual correlation patterns for each of the expression profiles of each of the 67 genes separately, by computing a 67×67 gene-to-gene correlation matrix, with the level of correlation between two genes taken as the Pearson correlation coefficient between their cortical expression profiles. This matrix was subdivided into clusters (also called modules) using Newman’s modularity algorithm^57^, maximizing for each cluster the intra-cluster cohesion and minimizing cohesion between genes of different clusters. Per cluster, a cluster-level expression profile was computed by calculating the average gene expression profile across all genes in the given cluster, resulting in clusters of genes showing distinct brain maps.

#### Interaction gene-set analyses

Aiming to determine whether genes associated to both BV and intelligence are over- (or under-) expressed in specific brain regions, we performed interaction analysis as implemented in MAGMA (v1.07b). This analysis tested whether the combined involvement of a gene set (here: genes related to BV) and continuous gene properties (here: gene expression) is different from their individual effects on intelligence. A positive interaction effect would suggest that, within the set of BV-related genes, the relation between expression and gene-based Z-scores for intelligence is stronger compared to other genes (i.e., all genes for which both expression data from AHBA, as well as gene-based Z-scores for intelligence were available). This would indicate that gene expression in that specific region plays a particular role in the genetic relation between brain volume and intelligence.

An interaction analysis was conducted separately for gene expression in each of 57 brain regions defined in the Allen Human Brain Atlas^33^ (AHBA). To control for potential variation in general expression levels across genes, we conditioned on the marginal effects of the BV-related gene set and average gene expression across all 57 regions, as well as for their interaction. The set of BV-related genes consisted of all genes that were significant in MAGMA’s gene-based analysis and/or mapped from SNP-level results in FUMA. Interaction effects were deemed significant if their *P*-value exceeded the Bonferroni-corrected threshold of *P* < 8.77×10^−4^ (0.05/57).

## Data availability

Our policy is to make genome-wide summary statistics (sumstats) publicly available. Summary statistics from our brain volume meta-analyses are available for download at the website of the department of Complex Trait Genetics, CNCR (see **URLs**). The GWAS summary statistics from the intelligence meta-analysis conducted by Savage et al. (2018)^2^ are also available from this website.

## Acknowledgements

The full GWAS summary statistics from the ENIGMA-CHARGE collaboration were downloaded from http://enigma.ini.usc.edu/research/download-enigma-gwas-results/.

Data on infant head circumference has been contributed by EGG Consortium and was downloaded from www.egg-consortium.org.

The Genotype-Tissue Expression (GTEx) Project was supported by the Common Fund of the Office of the Director of the National Institutes of Health, and by NCI, NHGRI, NHLBI, NIDA, NIMH, and NINDS. The data used for the analyses described in this manuscript were obtained from the GTEx Portal (see **URLs**) on 06/12/2018.

This work was funded by The Netherlands Organization for Scientific Research (NWO Brain & Cognition 433-09-228, NWO MagW VIDI 452-12-014, NWO VICI 435-14-005 and 453-07-001, 645-000-003). P.R.J. was funded by the Sophia Foundation for Scientific Research (SSWO, grant nr: S14-27).

Analyses were carried out on the Genetic Cluster Computer, which is financed by the Netherlands Scientific Organization (NWO: 480-05-003), by the VU University, Amsterdam, the Netherlands, and by the Dutch Brain Foundation, and is hosted by the Dutch National Computing and Networking Services SurfSARA. This research has been conducted using the UK Biobank Resource (application number 16406). We would like to thank the participants and researchers who collected and contributed to the data.

## URLs

Early Growth Genetics consortium: www.egg-consortium.org

ENIGMA: http://enigma.ini.usc.edu/research/download-enigma-gwas-results/

FUMA GWAS platform: http://fuma.ctglab.nl/

LD Score regression: https://github.com/bulik/ldsc

MAGMA: https://ctg.cncr.nl/software/magma

MsigDB gene-sets: http://software.broadinstitute.org/gsea/msigdb/index.jsp

GTEx portal: https://www.gtexportal.org/home/

Mendelian randomization - GSMR: http://cnsgenomics.com/software/gsmr/

Multi-trait-based conditional & joint analysis using GWAS summary data (mtCOJO): http://cnsgenomics.com/software/gcta/#mtCOJO

COLOC R package: https://cran.r-project.org/web/packages/coloc/coloc.pdf

Allen Human Brain Atlas (AHBA): http://human.brain-map.org/

GeneCards: https://www.genecards.org/

## Supplementary Information includes

**1. Supplementary Methods**

1.1 UK Biobank

1.2 UK Biobank genotype data

**2. Supplementary Results**

1. Results of GWAS meta-analysis of BV

1.1 UKB, ENIGMA-CHARGE & EGG data

1.2 Functional annotation

1.3 Gene mapping results

1.4 Gene-set results

2. Validation GWAS meta-analysis

2.1 GWAS meta-analysis on strictly ICV in UKB and ENIGMA-CHARGE

2.2 Functional annotation

2.3 Gene mapping results

2.4 Gene-set results

2.5. Overlap between both meta-analyses

2.6. Gene-card summary of BV genes overlapping with intelligence

**3. Supplementary Figures (1 to 13)**

**4. Supplementary Tables (1 to 28)**

