## Supplementary Information & Figures for "GWAS of brain volume on 54,407 individuals and cross-trait analysis with intelligence identifies shared genomic loci and genes"

**Affiliations:**

**This PDF file includes:**

### SUPPLEMENTARY METHODS

#### 1.1 UK Biobank

We performed a genome-wide association study using data from the UK Biobank Study (UKB, [www.ukbiobank.ac.uk](http://www.ukbiobank.ac.uk)). The UKB is a prospective population-based cohort that collects a wide variety of data in over 500,000 participants throughout the UK^1^. The UKB study received ethical approval from the National Research Ethics Service Committee North West-Haydock (reference 11/NW/0382), and all study procedures were in accordance with the World Medical Association for medical research. Between 2006 and 2010, 9.2 million invitations were sent to potential participants between the age of 40 and 69 years who were living within 25 miles distance from one of the 22 UK Biobank research centers, resulting in N=503,325 individuals included at the start of the study. UK Biobank collects high dimensional data, including questionnaire data, anthropometric measurements, DNA collection and multi-organ magnetic resonance imaging (MRI). Our study included MRI imaging data of the brain, which was added to the study protocol from 2015 onwards, and second-release imputed genotype data of the full UKB cohort (available since March 2018).

#### 1.2 UK Biobank genotype data

This study used imputed genotype data of the full UKB population that were released in July 2017. Genotype data collection and processing was performed by the coordinating UKB team, and has been described previously^2^. In short, genotyping was carried out on DNA extracted from blood samples in 489,212 individuals. Two Affymetrix genotyping arrays we used with added custom content: 438,692 participants were genotyped on the UK Biobank Axiom array, and 50,520 participants on the UK Biobank BiLEVE Axiom array, which included 812,428 genetic markers. Genotype quality control selected 488,377 individuals and 805,426 markers. Imputation was performed by the UK Biobank coordinating team, and quality-controlled data were imputed to a combined reference panel that included the Haplotype Reference Consortium (version 1.1), the UK10K, and the 1000 Genomes project reference panels. Imputed genotype data were made available in 487,422 participants, which included 92,693,895 genetic variants in total.

The current study was performed in unrelated participants of European ancestry. Ancestry was determined by projecting 1000 Genomes project genetic principal components (PCs) on the UKB genotypes and ancestry was assigned based on the closest Mahalanobis distance from the 1000 Genomes project population average.

Genome-wide association analysis (GWAS) of intracranial volume (ICV) in the UKB cohort was carried out through linear regression in PLINK^3^. Imputed variants were converted to hard-call genotypes prior to performing GWAS using a certainty threshold of 0.9. Genetic analyses were corrected for age, sex, height, genotype array, UKB assessment center, Townsend deprivation index (TDI), and 10 genetic PCs. These PCs were calculated using FlashPCA^4^ in the subset of European individuals on a set of 145,432 genotyped common variants (MAF>0.01) that were linkage disequilibrium (LD) independent SNPs (*r^2^*<0.1). Post-imputation filtering consisted of imputation quality (INFO score <0.9) and low allele frequency (MAF<0.005).

### SUPPLEMENTARY RESULTS

#### 1. Results of GWAS meta-analysis of BV

##### 1.1 UKB, ENIGMA-CHARGE & EGG data

The main meta-analysis in the current study combined ICV GWAS results from the UK Biobank (UKB; N=17,062) with ICV data from the ENIGMA-CHARGE collaboration (N=26,577) and data on head circumference from the EGG consortium (N=10,678), resulting in a final meta-analysis of brain volume in N=54,407 individuals (**Supplementary Fig. 1**).

In order to assess whether there was inflation in the observed test statistics compared to expected values, we inspected the Lambda GC (λ_GC_) and the LDSC intercept of the meta-analytic results (**Supplementary Table 2**). A λ_GC_ >1 suggests an inflation of the genetic effects, which can be due to both spurious and genuine effects. The value of λ_GC_ is likely to increase with sample size^5^. An LDSC intercept >1 suggests that there is spurious association, although an intercept <1.10 is usually considered to suggest that the signal is mostly due to genuine association effects^6^. For the current meta-analysis, the λ_GC_ was 1.23, indicating that, compared to the expected distribution, the median test statistic was inflated. The LDSC intercept of 1.037 suggests that the observed inflation is mostly due to genuine polygenicity, rather than caused by population stratification. Lead SNPs across all 35 independent genomic risk loci had the same direction of effect in all cohorts in which they were available (**Supplementary Table 3**), thereby extending our confidence in the results.

##### 1.2 Functional annotation

To extract additional biological information from the GWAS results, we ran functional annotation analyses using the FUMA^7^ annotation platform. Annotation was performed on all SNPs that were considered to be a *candidate* SNP given high LD with one of the independent significant SNPs (*r^2^*>0.6) and a low *P-*value (*P*<1×10^-5^), which resulted in 5,802 *candidate* SNPs (**Supplementary Table 5**). We observed that most of these SNPs were located within intergenic regions (n=1,131, 19.5%) or intronic regions (n=2,879, 49.6%), and a small minority of SNPs in exonic regions (n=64, 1.1%; **Supplementary Fig. 5**). Compared to the functional categories of all SNPs included in the meta-analysis, we observed a higher proportion of candidate SNPs in intronic regions (49.6% vs. 37.5% overall, *P*=1.63×10^-81^), a lower proportion of intergenic SNPs (19.5% vs. 45.6% overall, *P*<1×10^-100^) and similar proportion of SNPs in exonic regions (1.1% vs 0.7% overall, *P*=0.001).

Of the SNPs annotated to exonic regions, there were 30 SNPs that had non-synonymous changes in the proteins of the gene (ExNS; **Supplementary Table 6**). These ExNS SNPs were located in exons of 15 genes, with several genes including multiple ExNS SNPs: the *SPPL2C* gene contained 8 ExNS SNPs, all located in exon 1 of this gene. This gene codes for the signal peptide peptidase-like 2C, an intramembrane protease (IMP) that has been shown to play a role in the degradation of signaling peptides in the brain^8^, of which its exact biological substrate remains largely unclear^9^, but has recently been suggested to protease SNARE proteins^10^. In addition, the *MAPT* gene contained 5 ExNS SNPs in two exons, of which 4 in exon 6 and 1 in exon 8. In addition, *MAPT* contained 1 stop-gain SNP, located in exon 1. The *MAPT* gene product is the Microtubule Associated Protein Tau protein, a protein that regulates the assembly of microtubules^11^, and which has been linked to a wide variety of neurodegenerative disorders including Alzheimer’s disease^12^, frontotemporal dementia^13^ and Parkinson’s disease^14^.

##### 1.3 Gene mapping results

To find genes for brain volumes, we used three strategies to map SNPs to genes in FUMA^15^, and genome-wide gene-based association analysis (GWGAS) conducted in MAGMA^16^ (**Online Methods**; **Supplementary Fig. 6**). Summary statistics of the meta-analysis of brain volume (UKB, ENIGMA-CHARGE, EGG) were used as input for FUMA. FUMA gene-mapping mapped the 35 genomic loci to 101 genes through positional mapping (locus near to, or within a gene), 187 genes through eQTL mapping (locus associated with the expression of a gene), and 201 genes through chromatin-chromatin interactions (locus that physically interacts with a close or distant gene through the 3D structure of the genome), leading to a set of 340 unique genes (**Fig 2a**; **Supplementary Table 8)**. Of these genes, 111 genes (32.6%) were mapped by more than one gene-mapping method and 37 genes (10.9%) by all three mapping methods. Of all mapped genes associated with BV in FUMA, 73 genes had a pLI > 0.90, suggesting extreme intolerance to loss of function (LoF) mutations (**Supplementary Table 8**). In addition, genome-wide gene-based analysis (GWGAS) was carried out in MAGMA^16^ . These analyses implicated a role of 70 significant genes (**Fig. 2b**; **Supplementary Table 9**), of which a large part (11 genes, 15.7%) were mapped on chromosome 17. Of the 70 significant genes, 47 genes (67.1%) were also observed in FUMA and 16 genes (22.9%) were implicated through all three mapping strategies (positional mapping, eQTL mapping and chromatin-chromatin interaction mapping; *FRZB, FOXO3, CDK6, FAM49B, INA, PTEN, WBP1L, ERBB3, HMGA2, MAP3K12, PRR13, RAB5B, AJUBA, HAUS4, PRMT5, RBM23*). Of these 16 genes that were observed in all gene-mapping methods, several genes are known to be important in several regulatory functions of cell-signaling and cell neuronal development. *CKD6* codes the cyclin dependent kinase 6 protein, involved in G1 phase cell-cycle regulation through the phosphorylation of the pRB protein^17^. The *FOXO3* gene is part of the forkhead gene-family that code for transcription factors involved in cell survival. Transcription factors coded by *FOXO3* are phosphorylated by *AKT1* ^18^ and induce cell death through triggering of apoptosis. Overexpression of this gene has been shown to lead to cell growth decline in cell lines^19^. *ERBB3* (also known as *HER3*: human epidermal growth factor receptor 3) is part of the epidermal growth factor receptor tyrosine kinases and is a receptor for the heregulin protein^20^. Activation of the ErBB3 receptor leads to activation of the PI3K/ATK-signaling pathway which modifies cell differentiation and proliferation^21^.

##### 1.4 Gene-set results

We performed gene-set analysis in MAGMA^16^, using *P*-values of 18,161 genes as input from prior gene-based testing, and testing 7,246 gene-sets extracted from the online molecular signatures database (MsigDB, version 6.0;) repository^22^. Selected gene sets included expert-curated gene sets, motif, computational, gene ontology (GO), oncogenic, and immunologic gene sets. We were able to identify 23 significant gene-sets (**Supplementary Table 12**). Almost all gene-sets showed partial overlap in genes with each other (**Supplementary Fig. 7**), and thus several significantly associated genes were shared between those gene-sets **(Supplementary Fig. 8**). Among these genes were *PTEN* (*n_genesets_*=13, *P*=3.70×10^-9^), *AKT3* (*n_genesets_*=14, *P*=5.12×10^-11^), and *FOXO3* (*n_genesets_*=14, *P*=6.54×10^-14^). The *PTEN* gene (Phosphatase And Tensin Homolog deleted on chromosome ten) plays an important role in controlling the phosphoinositide 3‐kinase signaling pathway involved in cell proliferation and cell survival regulation by dephosphorylation of phosphatidylinositol 3,4,5‐triphosphate^23^, and has shown to be an important tumor suppressor in several hereditary cancer syndromes, most importantly Cowden syndrome^24^. *AKT3,* a gene that codes for the Serine/Threonine Kinase 3, acts downstream on the PI3K pathway, and is, similar to the aforementioned *PTEN* gene^25^, involved in regulations of several cell processes, including cell proliferation and cell death^26^. *AKT3* has a prominent role in post-natal brain development^27,28^, and abnormal functioning of this gene is associated with abnormalities in brain development in humans^29^. Also, mutations in this gene have been linked to the development of several cancer types^30,31^. *FOXO3,* forkhead box O3 (see also **Supplementary Results 1.3**), is a transcription factor and a key factor in a wide array of cellular functions, including cell proliferation, differentiation and survival^32^. These results suggest that similar regulatory biological pathways are involved in normal cell division during brain development as well as in aberrant cell division in the development of cancerous tumors in several types of cancer, which explains why we observed significant enrichment of genes in small-cell lung cancer gene-sets.

#### 2. Validation GWAS meta-analysis

##### 2.1 GWAS meta-analysis on strictly ICV in UKB and ENIGMA-CHARGE

Since our main meta-analysis included 1) data on head circumference (which is a proven proxy^33–35^ of brain volume, but may introduce some heterogeneity in the genetic signal) and 2) data that was corrected for height on a post-hoc procedure using mtCOJO^36^, we here provide results of a second meta-analysis using a stricter phenotype definition of ICV in a subset of the data. This second meta-analysis combined ICV GWAS results from the UK Biobank (UKB; N=17,062 individuals) with ICV data from a subset of the ENIGMA-CHARGE collaboration for which height-adjusted GWAS summary statistics were available (N=21,875 individuals), resulting in a final sample size of N=38,937 individuals (**Supplementary Fig. 1**). This subset concerns individuals of whom height data were available to include in the genome-wide analysis as a covariate. In contrast to the Early Growth Genetics (EGG) consortium data, both UKB and ENIGMA-CHARGE use a measure of intracranial volume obtained through magnetic resonance imaging (MRI; T1-weighted). The genetic correlation between both cohorts, computed using LDSC^6,37^, did not deviate from 1 (*r_g_* = 1.01, SE = 0.08; **Supplementary Table 2**).

In order to assess whether there was inflation in the observed test statistics compared to expected values, we inspected the Lambda GC (λ_GC_) and the LDSC intercept of the meta-analytic results. For the meta-analysis of the UKB and the height-adjusted subset of ENIGMA-CHARGE the λ_GC_ was 1.17, indicating that the median test statistic was inflated compared to the expected distribution. The LDSC intercept of 1.027 suggests that the observed inflation is mostly due to genuine polygenicity, rather than caused by population stratification. All of the lead SNPs in the 25 genomic loci identified in the meta-analysis had the same direction of effect in both cohorts (**Supplementary Table 15**), thereby extending our confidence in the results.

##### 2.2 Functional annotation

Analogous to the primary meta-analysis, annotation was performed on all SNPs that were considered to be a *candidate* SNP given high LD with one of the independent significant SNPs (*r^2^*>0.6) and a low *P-*value (*P* < 1×10^-5^), which resulted in 4,523 GWS SNPs (**Supplementary Table 16**) and 4,129 *candidate* SNPs (**Supplementary Table 17**). In line with the annotation based on the main meta-analysis, we observed that most of these SNPs were located within intergenic regions (n=641, 14.2%) or intronic regions (n=2,102, 46.5%), and a minority of SNPs in exonic regions (n=55, 1.2%). By testing for heritability enrichment of SNP categories with partitioned LD Score regression, we found significant heritability enrichment in 9 SNP categories of which all but one had also been found using the GWAS meta-analysis of BV (**Supplementary Table 18**). Interestingly, we found significant enrichment of SNPs located in promotor-flanking regions (enrichment=14.8, SE=4.3, *P*=1.36×10^-3^) had a low *P-*value in the meta-analysis of BV but did not pass the multiple testing corrected threshold in the primary meta-analysis (enrichment=10.4, SE=4.4, *P=*0.03).

Of the SNPs annotated to exonic regions, there were 27 SNPs categorized that had non-synonymous changes in the proteins of the gene (**Supplementary Table 19**). These ExNS SNPs were located in exons of 12 genes, with five genes containing multiple ExNS SNPs (*CRHR1, KANSL1, MAPT, PDCD11, SPPL2C*). The *CRHR1* gene is a protein-coding gene that codes for the Corticotropin Releasing Hormone Receptor 1^38^, and is highly expressed in several brain areas including hippocampus^39^, and plays a regulatory role in response to stress^40^.

The *MAPT* gene contained 5 ExNS SNPs in two exons, of which 4 in exon 6 and 1 in exon 8. In addition, *MAPT* contained one stop-gain SNP, located in exon 1. This gene codes for the microtubule-associated protein tau. Aberrant microtubule assembly by dysfunctioning tau protein is closely linked to the risk of neurodegenerative disorders such as Alzheimer’s disease^41^ and Parkinson’s disease^42^. The *PDCD11* gene (programmed cell-death 11) is an apoptosis-related gene located on chromosome 10, which contained two annotated ExNS in codon 24 and 36. The protein coded by this gene functions as a cell-death regulator and is able to induce apoptosis through the transcription of transmembrane fas ligands^43^.

##### 2.3 Gene mapping results

Summary statistics of the second meta-analysis (UKB and height-adjusted subset of ENIGMA-CHARGE) were used as input for FUMA and MAGMA. Positional, eQTL, and chromatin interaction (CI) mapping of the SNPs in the genomic risk loci resulted in 64, 96 and 119 genes, respectively, whereas GWGAS identified 71 genes (**Supplementary Tables 20, 21**). Of the 12 genes identified using all 4 methods (*PRMT5*, *FRZB*, *FAM49B*, *PRR13*, *WBP1L*, *HAUS4*, *RBM23*, *PTEN*, *MAP3K12*, *HMGA2*, *AJUBA*, *INA*), *HMGA2* had the lowest gene-based *P*-value (*P* = 1.54×10^-13^). The *HMGA2* gene is an oncogene that is often involved in tumorigenesis^44^ and plays a role in cell-differentiation^45^ and cell-aging^46^ in normal development. Of all genes that were associated to ICV, 44 had a pLI > 0.90, suggesting extreme intolerance to loss of function mutations

##### 2.4 Gene-set results

We performed gene-set analysis in MAGMA^16^, using *P*-values of 18,094 genes as input from prior gene-based testing, and testing 7,426 gene-sets extracted from the online molecular signatures database (MsigDB, version 6.0;) repository^22^. Selected gene sets included expert curated gene sets, motif, computational, gene ontology (GO), oncogenic, and immunologic gene sets. We were able to identify 16 significant gene-sets (**Supplementary Table 22**). The top most significant gene-sets were the *BIOCARTA: Longevity pathway* (*P=*1.73×10^-9^), *REACTOME: PI3K events in Erbb2 signaling* (*P=*2.45×10^-8^) and *GO: Central nervous system neuron axogenesis* (*P=*7.76×10^-8^), all of which had been observed in the meta-analysis of BV (**Supplementary Table 12**). In addition, there were several gene-sets that had not been associated with BV but were significant in the analysis of ICV, including *GO: telencephalon development* (*P*=3.96×10^-6^), *GO: insulin-like growth factor receptor signaling pathway* (*P*=5.57×10^-6^) and *GO: regulation of cellular response to stress* (*P=*4.30×10^-6^).

Next, we combined the MAGMA gene-based association testing with gene-expression data (RNA sequencing) from 53 tissue types from the GTEx consortium^47^, to find associations between gene-associations and the tissue-specific expression. The GTEx consortium database contains RNA sequencing data samples from 13 brain-tissue types. Using the gene-based *P-*values of BV from MAGMA association test, we did not find signification associations in tissue gene-expression (**Supplementary Table 22)**, potentially due to insufficient power of the GWAS. Alternatively, this negative finding may be explained by a lack of specificity of the genetic signal for one type of tissue.

To further investigate whether the genetic signal could be linked to gene-expression within single brain cell-types, we performed gene-expression analyses in MAGMA using RNA sequencing data from 565 brain cell-types derived from adult mouse brain^48^. After correction for testing multiple gene-expression patterns, none of these cell-types passed the Bonferroni threshold for significance (**Supplementary Table 22**). By ranking the single cell-types with the lowest *P-*value, several types of polydendrocyte were the most significant in the analyses, including Tnr-Pdgfa-Pik3r3-polydendrocytes (P=7.46×10^-4^), Tnr-Cspg5-polydendrocyte/NG2 cells (*P=*2.53×10^-3^), and Tnr-Bmp4-polydendrocyte (*P*=7.46×10^-4^) cells. Polydendrocytes of the brain fulfil a wide range of functions^49^, including myelin maintenance and repair^50^, and are precursor cell-types that can develop into oligodendrocytes, astrocytes and possibly also into neurons^51^.

##### 2.5 Overlap between both meta-analyses

As mentioned previously, a second meta-analysis was conducted in order to validate our initial findings. The strong genetic correlation between ICV in the UKB and ENIGMA-CHARGE samples and head circumference in the EGG data warrants meta-analysis. In addition, the high genetic correlation between the ENIGMA-CHARGE ICV data that was corrected for height using mtCOJO^36^ and the subset of the ENIGMA-CHARGE data for which height was included as a covariate in the original analyses (*r_g_*=0.997, SE=0.01; **Supplementary Table 2**) suggests that we successfully remove confounding effects of height by first creating height-conditioned GWAS results using mtCOJO. However, by comparing the results of both meta-analyses we hope to provide compelling evidence that our main meta-analytic approach is actually 1) tagging ICV signal and 2) providing increased power over using a stricter phenotype.

Compared to the validation GWAS meta-analysis, the primary meta-analysis expanded the sample size by adding (1) GWAS results of infant head circumference (HC) from the Early Growth Genetics (EGG) consortium^52^ (N=10,768) and (2) samples from the ENIGMA-CHARGE collaboration for which height was not available as a covariate. This resulted in a total sample size of N = 54,407 (i.e., an increase in sample size of N=15,470, i.e., of 39,7% compared to the height-adjusted GWAS).

Given the strong phenotypic correlation (*r*=0.55 in UK Biobank) between height and ICV (as well as HC), we consider it important to adjust for height in the analysis, so that the obtained genetic signal for brain volume is not contaminated by genetic signal for height. The LDSC-computed genetic correlation between the primary meta-analysis and the height-adjusted analysis approximated 1 (*r_g_*=0.96, SE=0.01), suggesting that most genetic signal was shared. Of the 25 loci that were identified in the validation meta-analysis, 23 overlapped with the 35 loci identified in the primary meta-analysis (**Supplementary Fig. 12**; **Supplementary Table 15)**. Out of these 23 loci, 9 had the same lead SNP across both meta-analyses. Gene-mapping results from MAGMA and FUMA identified 217 unique genes for the validation meta-analysis, of which only 12 were not shared with the primary meta-analysis (**Supplementary Fig. 12**; **Supplementary Tables 20, 21**). Gene-set analysis based on the MAGMA gene-based *P-*values for the validation meta-analysis showed 16 significant gene sets (**Supplementary Table 22**), of which 13 were also observed in the height-adjusted analysis and 3 were different. We did not observe any significant tissue type expression in the gene-expression analyses.

##### 2.6 Gene-card summary of BV genes overlapping with intelligence

Gene-mapping strategies in FUMA and MAGMA identified 67 overlapping genes between the BV GWAS meta-analysis and the GWAS meta-analysis of intelligence (**Supplementary Table 24**). Below we provide a description of these genes using GeneCard^53^ summaries.

*1. AMBRA1*

**GeneCards:** AMBRA1 (Autophagy And Beclin 1 Regulator 1) is a Protein Coding gene. Diseases associated with AMBRA1 include Selective Immunoglobulin Deficiency Disease and Central Retinal Artery Occlusion. Among its related pathways are Autophagy Pathway and Neuroscience. Gene Ontology (GO) annotations related to this gene include ubiquitin protein ligase binding.

*2. ARHGAP27*

**GeneCards:** ARHGAP27 (Rho GTPase Activating Protein 27) is a Protein Coding gene. Among its related pathways are p75 NTR receptor-mediated signaling and Signaling by Rho GTPases. Gene Ontology (GO) annotations related to this gene include GTPase activator activity and SH3 domain binding. An important paralog of this gene is ARHGAP12.

**Entrez Gene Summary:** This gene encodes a member of a large family of proteins that activate Rho-type guanosine triphosphate (GTP) metabolizing enzymes. The encoded protein may pay a role in clathrin-mediated endocytosis. Alternatively spliced transcript variants encoding multiple isoforms have been observed for this gene. [provided by RefSeq, Aug 2013]

*3. ARIH2*

**Entrez Gene Summary:** The protein encoded by this gene is an E3 ubiquitin-protein ligase that polyubiquitinates some proteins, tagging them for degradation. The encoded protein upregulates p53 in some cancer cells and may inhibit myelopoiesis. Several transcript variants encoding different isoforms have been found for this gene, although the full-length nature of some of them have not been determined yet. [provided by RefSeq, Nov 2015]

**GeneCards:** ARIH2 (Ariadne RBR E3 Ubiquitin Protein Ligase 2) is a Protein Coding gene. Among its related pathways are Class I MHC mediated antigen processing and presentation and NF-kappaB Signaling. Gene Ontology (GO) annotations related to this gene include nucleic acid binding and ubiquitin-protein transferase activity.

*4. ARL17A*

**Entrez Gene Summary:** ARL17A (ADP Ribosylation Factor Like GTPase 17A) is a Protein Coding gene. Gene Ontology (GO) annotations related to this gene include *GTP binding*. An important paralog of this gene is ARL17B.

*5. ARL17B*

**GeneCards:** ARL17B (ADP Ribosylation Factor Like GTPase 17B) is a Protein Coding gene. Gene Ontology (GO) annotations related to this gene include *GTP binding*. An important paralog of this gene is ARL17A.

*6. ARMC2*

**GeneCards:** ARMC2 (Armadillo Repeat Containing 2) is a Protein Coding gene. Gene Ontology (GO) annotations related to this gene include binding.

*7. AS3MT*

**Entrez Gene Summary:** AS3MT catalyzes the transfer of a methyl group from S-adenosyl-L-methionine (AdoMet) to trivalent arsenical and may play a role in arsenic metabolism (Lin et al., 2002 [PubMed 11790780]).[supplied by OMIM, Mar 2008]

**GeneCards:** AS3MT (Arsenite Methyltransferase) is a Protein Coding gene. Diseases associated with AS3MT include Griscelli Syndrome, Type 3. Among its related pathways are Cytochrome P450 - arranged by substrate type and arsenate detoxification I (glutaredoxin). Gene Ontology (GO) annotations related to this gene include *methyltransferase activity* and *protein-L-isoaspartate (D-aspartate) O-methyltransferase activity*.

*8. ATG13*

**Entrez Gene Summary:** The protein encoded by this gene is an autophagy factor and a target of the TOR kinase signaling pathway. The encoded protein is essential for autophagosome formation and mitophagy. [provided by RefSeq, Oct 2016]

**GeneCards:** ATG13 (Autophagy Related 13) is a Protein Coding gene. Among its related pathways are PI3K-AKT-mTOR signaling pathway and therapeutic opportunitiesand Autophagy Pathway. Gene Ontology (GO) annotations related to this gene include *protein kinase binding*.

*9. AVL9*

**GeneCards:** AVL9 (AVL9 Cell Migration Associated) is a Protein Coding gene

*10. C10orf32*

**Entrez Gene Summary:** This locus represents naturally occurring read-through transcription between the neighboring **C10orf32** (chromosome 10 open reading frame 32) and AS3MT (arsenic, +3 oxidation state, methyltransferase) genes. The read-through transcript is a candidate for nonsense-mediated mRNA decay (NMD), and is therefore unlikely to produce a protein product. [provided by RefSeq, Dec 2010]

**GeneCards:** BORCS7-ASMT (BORCS7-ASMT Readthrough (NMD Candidate)) is an RNA Gene, and is affiliated with the ncRNA class. An important paralog of this gene is BORCS7.

*11. C8orf82*

**GeneCards:** C8orf82 (Chromosome 8 Open Reading Frame 82) is a Protein Coding gene.

*12. CAMTA1*

**Entrez Gene Summary:** The protein encoded by this gene contains a CG1 DNA-binding domain, a transcription factor immunoglobulin domain, ankyrin repeats, and calmodulin-binding IQ motifs. The encoded protein is thought to be a transcription factor and may be a tumor suppressor. However, a translocation event is sometimes observed between this gene and the WWTR1 gene, with the resulting WWTR1-CAMTA1 oncoprotein leading to epithelioid hemangioendothelioma, a malignant vascular cancer. [provided by RefSeq, Mar 2017]

**GeneCards:** CAMTA1 (Calmodulin Binding Transcription Activator 1) is a Protein Coding gene. Diseases associated with CAMTA1 include Cerebellar Ataxia, Nonprogressive, With Mental Retardationand Epithelioid Hemangioendothelioma.

*13. CCHCR1*

**Entrez Gene Summary:** This gene encodes a protein with five coiled-coil alpha-helical rod domains that is thought to act as a regulator of mRNA metabolism through its interaction with mRNA-decapping protein 4. It localizes to P-bodies, the site of mRNA metabolism, with an N-terminus that is required for this subcellular localization, suggesting it is a P-body component. Naturally occurring mutations in this gene are associated with psoriasis. [provided by RefSeq, May 2017]

**GeneCards:** CCHCR1 (Coiled-Coil Alpha-Helical Rod Protein 1) is a Protein Coding gene. Diseases associated with CCHCR1 include Psoriasis and Peeling Skin Syndrome. An important paralog of this gene is CCDC57.

*14. CD164*

**Entrez Gene Summary: T**his gene encodes a transmembrane sialomucin and cell adhesion molecule that regulates the proliferation, adhesion and migration of hematopoietic progenitor cells. The encoded protein also interacts with the C-X-C chemokine receptor type 4 and may regulate muscle development. Elevated expression of this gene has been observed in human patients with Sezary syndrome, a type of blood cancer, and a mutation in this gene may be associated with impaired hearing. [provided by RefSeq, Oct 2016]

**GeneCards:** CD164 (CD164 Molecule) is a Protein Coding gene. Diseases associated with CD164 include Deafness, Autosomal Dominant 66 and Pollen Allergy. Among its related pathways are Hematopoietic Stem Cell Differentiation Pathways and Lineage-specific Markers and Lysosome. An important paralog of this gene is CD164L2.

*15. CENPW*

**GeneCards:** CENPW (Centromere Protein W) is a Protein Coding gene. Among its related pathways are Cell Cycle, Mitotic and Chromosome Maintenance. Gene Ontology (GO) annotations related to this gene include *protein heterodimerization activity*.

*16. CEP57L1*

**GeneCards:** CEP57L1 (Centrosomal Protein 57 Like 1) is a Protein Coding gene. Gene Ontology (GO) annotations related to this gene include *identical protein binding*and *gamma-tubulin binding*. An important paralog of this gene is CEP57.

*17. CRHR1*

**Entrez Gene Summary:** This gene encodes a G-protein coupled receptor that binds neuropeptides of the corticotropin releasing hormone family that are major regulators of the hypothalamic-pituitary-adrenal pathway. The encoded protein is essential for the activation of signal transduction pathways that regulate diverse physiological processes including stress, reproduction, immune response and obesity. Alternative splicing results in multiple transcript variants. Naturally-occurring readthrough transcription between this gene and upstream GeneID:147081 results in transcripts that encode isoforms that share similarity with the products of this gene. [provided by RefSeq, Aug 2016]

**GeneCards:** CRHR1 (Corticotropin Releasing Hormone Receptor 1) is a Protein Coding gene. Diseases associated with CRHR1 include Irritable Bowel Syndrome and Depression. Among its related pathways are Presynaptic function of Kainate receptors and G alpha (s) signalling events. Gene Ontology (GO) annotations related to this gene include *G protein-coupled receptor activity*. An important paralog of this gene is LINC02210-CRHR1.

*18. DALRD3*

**Entrez Gene Summary:** The exact function of this gene is not known. It encodes a protein with a DALR anticodon binding domain similar to that of class Ia aminoacyl tRNA synthetases. This gene is located in a cluster of genes (with a complex sense-anti-sense genome architecture) on chromosome 3, and contains two micro RNA (miRNA) precursors (mir-425 and mir-191) in one of its introns. Preferential expression of this gene (the miRNAs and other genes in the cluster) in testis suggests a role of this gene in spermatogenesis (PMID:19906709). [provided by RefSeq, Feb 2013]

**GeneCards:** DALRD3 (DALR Anticodon Binding Domain Containing 3) is a Protein Coding gene. Diseases associated with DALRD3 include Mitochondrial Complex I Deficiency. Gene Ontology (GO) annotations related to this gene include *nucleotide binding* and *arginine-tRNA ligase activity*.

*19. DCAKD*

**GeneCards:** DCAKD (Dephospho-CoA Kinase Domain Containing) is a Protein Coding gene. Among its related pathways are Integrated Breast Cancer Pathway. Gene Ontology (GO) annotations related to this gene include *dephospho-CoA kinase activity*.

*20. DYRK1A*

**Entrez Gene Summary:** This gene encodes a member of the Dual-specificity tyrosine phosphorylation-regulated kinase (DYRK) family. This member contains a nuclear targeting signal sequence, a protein kinase domain, a leucine zipper motif, and a highly conservative 13-consecutive-histidine repeat. It catalyzes its autophosphorylation on serine/threonine and tyrosine residues. It may play a significant role in a signaling pathway regulating cell proliferation and may be involved in brain development. This gene is a homolog of Drosophila mnb (minibrain) gene and rat Dyrk gene. It is localized in the Down syndrome critical region of chromosome 21, and is considered to be a strong candidate gene for learning defects associated with Down syndrome. Alternative splicing of this gene generates several transcript variants differing from each other either in the 5' UTR or in the 3' coding region. These variants encode at least five different isoforms. [provided by RefSeq, Jul 2008]

**GeneCards:** DYRK1A (Dual Specificity Tyrosine Phosphorylation Regulated Kinase 1A) is a Protein Coding gene. Diseases associated with DYRK1A include Mental Retardation, Autosomal Dominant 7 and Intellectual Disability Syndrome Due To A Dyrk1a Point Mutation. Among its related pathways are Cell Cycle, Mitoticand Regulation of lipid metabolism Insulin signaling-generic cascades. Gene Ontology (GO) annotations related to this gene include *identical protein binding*and *protein kinase activity*.

*21. EFCAB5*

**GeneCards:** EFCAB5 (EF-Hand Calcium Binding Domain 5) is a Protein Coding gene. Gene Ontology (GO) annotations related to this gene include *calcium ion binding*. An important paralog of this gene is NSRP1.

*22. EFTUD2*

**Entrez Gene Summary:** This gene encodes a GTPase which is a component of the spliceosome complex which processes precursor mRNAs to produce mature mRNAs. Mutations in this gene are associated with mandibulofacial dysostosis with microcephaly. Multiple transcript variants encoding different isoforms have been found for this gene. [provided by RefSeq, Apr 2012]

**GeneCards:** EFTUD2 (Elongation Factor Tu GTP Binding Domain Containing 2) is a Protein Coding gene. Diseases associated with EFTUD2 include Mandibulofacial Dysostosis, Guion-Almeida Type and Dysostosis. Among its related pathways are mRNA Splicing - Major Pathway and mRNA Splicing - Minor Pathway. Gene Ontology (GO) annotations related to this gene include *GTPase activity*.

*23. ERBB3*

**Entrez Gene Summary:** This gene encodes a member of the epidermal growth factor receptor (EGFR) family of receptor tyrosine kinases. This membrane-bound protein has a neuregulin binding domain but not an active kinase domain. It therefore can bind this ligand but not convey the signal into the cell through protein phosphorylation. However, it does form heterodimers with other EGF receptor family members which do have kinase activity. Heterodimerization leads to the activation of pathways which lead to cell proliferation or differentiation. Amplification of this gene and/or overexpression of its protein have been reported in numerous cancers, including prostate, bladder, and breast tumors. Alternate transcriptional splice variants encoding different isoforms have been characterized. One isoform lacks the intermembrane region and is secreted outside the cell. This form acts to modulate the activity of the membrane-bound form. Additional splice variants have also been reported, but they have not been thoroughly characterized. [provided by RefSeq, Jul 2008]

**GeneCards:** ERBB3 (Erb-B2 Receptor Tyrosine Kinase 3) is a Protein Coding gene. Diseases associated with ERBB3 include Lethal Congenital Contracture Syndrome 2and Retroperitoneal Leiomyosarcoma. Among its related pathways are GPCR Pathway and NFAT and Cardiac Hypertrophy. Gene Ontology (GO) annotations related to this gene include *protein homodimerization activity* and *transferase activity, transferring phosphorus-containing groups*.

*24. FOXO3*

**Entrez Gene Summary:** This gene belongs to the forkhead family of transcription factors which are characterized by a distinct forkhead domain. This gene likely functions as a trigger for apoptosis through expression of genes necessary for cell death. Translocation of this gene with the MLL gene is associated with secondary acute leukemia. Alternatively spliced transcript variants encoding the same protein have been observed. [provided by RefSeq, Jul 2008]

**GeneCards:** FOXO3 (Forkhead Box O3) is a Protein Coding gene. Diseases associated with FOXO3 include [Chromosome 6Q Deletion](http://www.malacards.org/card/chromosome_6q_deletion) and [Rhabdomyosarcoma](http://www.malacards.org/card/rhabdomyosarcoma). Among its related pathways are [PI3K-AKT-mTOR signaling pathway and therapeutic opportunities](http://pathcards.genecards.org/card/pi3k-akt-mtor_signaling_pathway_and_therapeutic_opportunities) and [Kit receptor signaling pathway](http://pathcards.genecards.org/card/kit_receptor_signaling_pathway). Gene Ontology (GO) annotations related to this gene include *DNA-binding transcription factor activity* and *protein kinase binding*.

*25. FZD2*

**Entrez Gene Summary:** This intronless gene is a member of the frizzled gene family. Members of this family encode seven-transmembrane domain proteins that are receptors for the wingless type MMTV integration site family of signaling proteins. This gene encodes a protein that is coupled to the beta-catenin canonical signaling pathway. Competition between the wingless-type MMTV integration site family, member 3A and wingless-type MMTV integration site family, member 5A gene products for binding of this protein is thought to regulate the beta-catenin-dependent and -independent pathways. [provided by RefSeq, Dec 2010]

**GeneCards:** FZD2 (Frizzled Class Receptor 2) is a Protein Coding gene. Diseases associated with FZD2 include Omodysplasia 2 and Autosomal Dominant Robinow Syndrome. Among its related pathways are Hippo signaling pathway and Nanog in Mammalian ESC Pluripotency. Gene Ontology (GO) annotations related to this gene include *G protein-coupled receptor activity* and *PDZ domain binding*.

*26. GBF1*

**Entrez Gene Summary:** This gene encodes a member of the Sec7 domain family. The encoded protein is a guanine nucleotide exchange factor that regulates the recruitment of proteins to membranes by mediating GDP to GTP exchange. The encoded protein is localized to the Golgi apparatus and plays a role in vesicular trafficking by activating ADP ribosylation factor 1. The encoded protein has also been identified as an important host factor for viral replication. Multiple transcript variants have been observed for this gene. [provided by RefSeq, Dec 2010]

**GeneCards:** GBF1 (Golgi Brefeldin A Resistant Guanine Nucleotide Exchange Factor 1) is a Protein Coding gene. Diseases associated with GBF1 include Legionnaires' Disease and Periventricular Nodular Heterotopia. Among its related pathways are Metabolism of proteins and Cargo trafficking to the periciliary membrane. Gene Ontology (GO) annotations related to this gene include *binding* and *ARF guanyl-nucleotide exchange factor activity*

*27. GCDH*

**Entrez Gene Summary:** The protein encoded by this gene belongs to the acyl-CoA dehydrogenase family. It catalyzes the oxidative decarboxylation of glutaryl-CoA to crotonyl-CoA and CO(2) in the degradative pathway of L-lysine, L-hydroxylysine, and L-tryptophan metabolism. It uses electron transfer flavoprotein as its electron acceptor. The enzyme exists in the mitochondrial matrix as a homotetramer of 45-kD subunits. Mutations in this gene result in the metabolic disorder glutaric aciduria type 1, which is also known as glutaric acidemia type I. Alternative splicing of this gene results in multiple transcript variants. A related pseudogene has been identified on chromosome 12. [provided by RefSeq, Mar 2013]

**GeneCards:** GCDH (Glutaryl-CoA Dehydrogenase) is a Protein Coding gene. Diseases associated with GCDH include Glutaric Acidemia I and Athetosis. Among its related pathways are Histidine, lysine, phenylalanine, tyrosine, proline and tryptophan catabolism and superpathway of tryptophan utilization. Gene Ontology (GO) annotations related to this gene include *flavin adenine dinucleotide binding* and *fatty-acyl-CoA binding*.

*28. GOSR1*

**Entrez Gene Summary:** This gene encodes a trafficking membrane protein which transports proteins among the endoplasmic reticulum and the Golgi and between Golgi compartments. This protein is considered an essential component of the Golgi SNAP receptor (SNARE) complex. Alternatively spliced transcript variants encoding distinct isoforms have been found for this gene. [provided by RefSeq, Jul 2008]

**GeneCards:** GOSR1 (Golgi SNAP Receptor Complex Member 1) is a Protein Coding gene. Diseases associated with GOSR1 include Chromosome 17Q11.2 Deletion Syndrome, 1.4-Mb. Among its related pathways are Metabolism of proteins and Vesicle-mediated transport. Gene Ontology (GO) annotations related to this gene include *SNAP receptor activity*.

*29. GOSR2*

**Entrez Gene Summary:** This gene encodes a trafficking membrane protein which transports proteins among the medial- and trans-Golgi compartments. Due to its chromosomal location and trafficking function, this gene may be involved in familial essential hypertension. [provided by RefSeq, Mar 2016]

**GeneCards:**
GOSR2 (Golgi SNAP Receptor Complex Member 2) is a Protein Coding gene. Diseases associated with GOSR2 include Epilepsy, Progressive Myoclonic, 6and Myoclonic Epilepsy Of Unverricht And Lundborg. Among its related pathways are Metabolism of proteins and Unfolded Protein Response (UPR). Gene Ontology (GO) annotations related to this gene include *transporter activity* and *SNAP receptor activity*. An important paralog of this gene is ENSG00000262633.

*30. GTF3C6*

**Entrez Gene Summary:** RNA polymerases are unable to initiate RNA synthesis in the absence of additional proteins called general transcription factors (GTFs). GTFs assemble in a complex on the DNA promoter and recruit the RNA polymerase. GTF3C family proteins (e.g., GTF3C1, MIM 603246) are essential for RNA polymerase III to make a number of small nuclear and cytoplasmic RNAs, including 5S RNA (MIM 180420), tRNA, and adenovirus-associated (VA) RNA of both cellular and viral origin.[supplied by OMIM, Mar 2008]

**GeneCards:** GTF3C6 (General Transcription Factor IIIC Subunit 6) is a Protein Coding gene. Among its related pathways are RNA Polymerase III Transcription Initiationand Activated PKN1 stimulates transcription of AR (androgen receptor) regulated genes KLK2 and KLK3.

*31. HLA-C*

**Entrez Gene Summary:** HLA-C belongs to the HLA class I heavy chain paralogues. This class I molecule is a heterodimer consisting of a heavy chain and a light chain (beta-2 microglobulin). The heavy chain is anchored in the membrane. Class I molecules play a central role in the immune system by presenting peptides derived from endoplasmic reticulum lumen. They are expressed in nearly all cells. The heavy chain is approximately 45 kDa and its gene contains 8 exons. Exon one encodes the leader peptide, exons 2 and 3 encode the alpha1 and alpha2 domain, which both bind the peptide, exon 4 encodes the alpha3 domain, exon 5 encodes the transmembrane region, and exons 6 and 7 encode the cytoplasmic tail. Polymorphisms within exon 2 and exon 3 are responsible for the peptide binding specificity of each class one molecule. Typing for these polymorphisms is routinely done for bone marrow and kidney transplantation. Over one hundred HLA-C alleles have been described [provided by RefSeq, Jul 2008]

**GeneCards:** HLA-C (Major Histocompatibility Complex, Class I, C) is a Protein Coding gene. Diseases associated with HLA-C include Psoriasis 1 and Human Immunodeficiency Virus Type 1. Among its related pathways are Cytokine Signaling in Immune system and Class I MHC mediated antigen processing and presentation. Gene Ontology (GO) annotations related to this gene include signaling receptor binding and TAP binding. An important paralog of this gene is HLA-B.

*32. HOOK2*

**Entrez Gene Summary:** Hook proteins are cytosolic coiled-coil proteins that contain conserved N-terminal domains, which attach to microtubules, and more divergent C-terminal domains, which mediate binding to organelles. The Drosophila Hook protein is a component of the endocytic compartment.[supplied by OMIM, Apr 2004]

**GeneCards:** HOOK2 (Hook Microtubule Tethering Protein 2) is a Protein Coding gene. Gene Ontology (GO) annotations related to this gene include identical protein binding.

*33. INA*

**Entrez Gene Summary:** Neurofilaments are type IV intermediate filament heteropolymers composed of light, medium, and heavy chains. Neurofilaments comprise the axoskeleton and they functionally maintain the neuronal caliber. They may also play a role in intracellular transport to axons and dendrites. This gene is a member of the intermediate filament family and is involved in the morphogenesis of neurons. [provided by RefSeq, Jun 2009]

**GeneCards:** INA (Internexin Neuronal Intermediate Filament Protein Alpha) is a Protein Coding gene. Diseases associated with INA include Wernicke Encephalopathy and Medulloepithelioma. Among its related pathways are Cytoskeleton remodeling Neurofilaments. Gene Ontology (GO) annotations related to this gene include *structural molecule activity* and *structural constituent of cytoskeleton*. An important paralog of this gene is NEFM.

*34. KANSL1*

**Entrez Gene Summary:** This gene encodes a nuclear protein that is a subunit of two protein complexes involved with histone acetylation, the MLL1 complex and the NSL1 complex. The corresponding protein in Drosophila interacts with K(lysine) acetyltransferase 8, which is also a subunit of both the MLL1 and NSL1 complexes. [provided by RefSeq, Jun 2012]

**GeneCards:** KANSL1 (KAT8 Regulatory NSL Complex Subunit 1) is a Protein Coding gene. Diseases associated with KANSL1 include Koolen-De Vries Syndrome and Koolen-De Vries Syndrome Due To A Point Mutation. Among its related pathways are Pathways Affected in Adenoid Cystic Carcinoma and Chromatin organization. Gene Ontology (GO) annotations related to this gene include *histone acetyltransferase activity (H4-K5 specific)* and *histone acetyltransferase activity (H4-K16 specific)*. An important paralog of this gene is KANSL1L.

*35. KIAA1919*

**Entrez Gene Summary:** MFSD4B (Major Facilitator Superfamily Domain Containing 4B) is a Protein Coding gene. Among its related pathways are Transport of glucose and other sugars, bile salts and organic acids, metal ions and amine compounds. An important paralog of this gene is MFSD4A.

*36. LRRC14*

**Entrez Gene Summary:** This gene encodes a leucine-rich repeat-containing protein. Alternate splicing results in multiple transcript variants. [provided by RefSeq, Dec 2012]

**GeneCards:** LRRC14 (Leucine Rich Repeat Containing 14) is a Protein Coding gene. Diseases associated with LRRC14 include Baller-Gerold Syndrome and Rapadilino Syndrome. An important paralog of this gene is LRRC14B.

*37. LRRC37A*

**GeneCards:** LRRC37A (Leucine Rich Repeat Containing 37A) is a Protein Coding gene. An important paralog of this gene is LRRC37A2.

*38. LRRC37A2*

**GeneCards:** LRRC37A2 (Leucine Rich Repeat Containing 37 Member A2) is a Protein Coding gene. An important paralog of this gene is LRRC37A.

*39. MAN2B1*

**Entrez Gene Summary:** This gene encodes an enzyme that hydrolyzes terminal, non-reducing alpha-D-mannose residues in alpha-D-mannosides. Its activity is necessary for the catabolism of N-linked carbohydrates released during glycoprotein turnover and it is member of family 38 of glycosyl hydrolases. The full length protein is processed in two steps. First, a 49 aa leader sequence is cleaved off and the remainder of the protein is processed into 3 peptides of 70 kDa, 42 kDa (D) and 13/15 kDa (E). Next, the 70 kDa peptide is further processed into three peptides (A, B and C). The A, B and C peptides are disulfide-linked. Defects in this gene have been associated with lysosomal alpha-mannosidosis. Alternatively spliced transcript variants encoding different isoforms have been found for this gene.[provided by RefSeq, Mar 2010]

**GeneCards:** MAN2B1 (Mannosidase Alpha Class 2B Member 1) is a Protein Coding gene. Diseases associated with MAN2B1 include Mannosidosis, Alpha B, Lysosomaland Alpha-Mannosidosis, Infantile Form. Among its related pathways are Lysosome and Innate Immune System. Gene Ontology (GO) annotations related to this gene include *carbohydrate binding* and *alpha-mannosidase activity*. An important paralog of this gene is MAN2B2.

*40. MAPT*

**Entrez Gene Summary:** This gene encodes the microtubule-associated protein tau (MAPT) whose transcript undergoes complex, regulated alternative splicing, giving rise to several mRNA species. MAPT transcripts are differentially expressed in the nervous system, depending on stage of neuronal maturation and neuron type. MAPT gene mutations have been associated with several neurodegenerative disorders such as Alzheimer's disease, Pick's disease, frontotemporal dementia, cortico-basal degeneration and progressive supranuclear palsy. [provided by RefSeq, Jul 2008]

**GeneCards:** MAPT (Microtubule Associated Protein Tau) is a Protein Coding gene. Diseases associated with MAPT include Frontotemporal Dementia and Supranuclear Palsy, Progressive, 1. Among its related pathways are Beta-Adrenergic Signaling and Kit receptor signaling pathway. Gene Ontology (GO) annotations related to this gene include *protein kinase binding* and *microtubule binding*.

*41. NSF*

**Entrez Gene Summary:** NSF (N-Ethylmaleimide Sensitive Factor, Vesicle Fusing ATPase) is a Protein Coding gene. Diseases associated with NSF include Tetanus and Neuronal Intranuclear Inclusion Disease. Among its related pathways are Metabolism of proteins and Golgi-to-ER retrograde transport. Gene Ontology (GO) annotations related to this gene include *protein kinase binding*.

*42. P4HTM*

**Entrez Gene Summary:** The product of this gene belongs to the family of prolyl 4-hydroxylases. This protein is a prolyl hydroxylase that may be involved in the degradation of hypoxia-inducible transcription factors under normoxia. It plays a role in adaptation to hypoxia and may be related to cellular oxygen sensing. Alternatively spliced variants encoding different isoforms have been identified. [provided by RefSeq, Jul 2008]

**GeneCards:** P4HTM (Prolyl 4-Hydroxylase, Transmembrane) is a Protein Coding gene. Diseases associated with P4HTM include Hypoxia. Gene Ontology (GO) annotations related to this gene include *calcium ion binding* and *iron ion binding*.

*43. PCBP2*

**Entrez Gene Summary:** The protein encoded by this gene appears to be multifunctional. Along with PCBP-1 and hnRNPK, it is one of the major cellular poly(rC)-binding proteins. The encoded protein contains three K-homologous (KH) domains which may be involved in RNA binding. Together with PCBP-1, this protein also functions as a translational coactivator of poliovirus RNA via a sequence-specific interaction with stem-loop IV of the IRES, promoting poliovirus RNA replication by binding to its 5'-terminal cloverleaf structure. It has also been implicated in translational control of the 15-lipoxygenase mRNA, human papillomavirus type 16 L2 mRNA, and hepatitis A virus RNA. The encoded protein is also suggested to play a part in formation of a sequence-specific alpha-globin mRNP complex which is associated with alpha-globin mRNA stability. This multiexon structural mRNA is thought to be retrotransposed to generate PCBP-1, an intronless gene with functions similar to that of PCBP2. This gene and PCBP-1 have paralogous genes (PCBP3 and PCBP4) which are thought to have arisen as a result of duplication events of entire genes. This gene also has two processed pseudogenes (PCBP2P1 and PCBP2P2). Multiple transcript variants encoding different isoforms have been found for this gene. [provided by RefSeq, Jan 2018]

**GeneCards:** PCBP2 (Poly(RC) Binding Protein 2) is a Protein Coding gene. Diseases associated with PCBP2 include Semantic Dementia and Leukemia, Chronic Myeloid. Among its related pathways are RIG-I/MDA5 mediated induction of IFN-alpha/beta pathways and mRNA Splicing - Major Pathway. Gene Ontology (GO) annotations related to this gene include *nucleic acid binding* and *RNA binding*.

*44. PLEKHM1*

**Entrez Gene Summary:** The protein encoded by this gene is essential for bone resorption, and may play a critical role in vesicular transport in the osteoclast. Mutations in this gene are associated with autosomal recessive osteopetrosis type 6 (OPTB6). Alternatively spliced transcript variants have been found for this gene. [provided by RefSeq, Sep 2009]

**GeneCards:** PLEKHM1 (Pleckstrin Homology And RUN Domain Containing M1) is a Protein Coding gene. Diseases associated with PLEKHM1 include Osteopetrosis, Autosomal Recessive 6 and Osteopetrosis, Autosomal Dominant 3.

*45. PPIL6*

**Entrez Gene Summary:** PPIL6 (Peptidylprolyl Isomerase Like 6) is a Protein Coding gene. Gene Ontology (GO) annotations related to this gene include peptidyl-prolyl cis-trans isomerase activity.

*46. PPP1R16A*

**Entrez Gene Summary:** Myosin light chain kinase and phosphatase (MLCP) complexes control the phosphorylation states of regulatory myosin light chains, which is crucial for muscle and intracellular movement. MLCPs typically contain a catalytic protein phosphatase 1 (PP1c) subunit, a myosin phosphatase targeting (MYPT) subunit, and another smaller subunit. The protein encoded by this gene represents an MYPT subunit, which is responsible for directing PP1c to its intended targets. However, while other MYPTs result in PP1c activation after becoming phosphorylated, the encoded protein is phosphorylated by protein kinase A and then inhibits the catalytic activity of PP1c. [provided by RefSeq, Jul 2016]

**GeneCards:** PPP1R16A (Protein Phosphatase 1 Regulatory Subunit 16A) is a Protein Coding gene. Among its related pathways are Beta-Adrenergic Signaling and Apoptotic Pathways in Synovial Fibroblasts. Gene Ontology (GO) annotations related to this gene include *actin binding*. An important paralog of this gene is PPP1R16B.

*47. PRKAR2A*

**Entrez Gene Summary:** cAMP is a signaling molecule important for a variety of cellular functions. cAMP exerts its effects by activating the cAMP-dependent protein kinase, which transduces the signal through phosphorylation of different target proteins. The inactive kinase holoenzyme is a tetramer composed of two regulatory and two catalytic subunits. cAMP causes the dissociation of the inactive holoenzyme into a dimer of regulatory subunits bound to four cAMP and two free monomeric catalytic subunits. Four different regulatory subunits and three catalytic subunits have been identified in humans. The protein encoded by this gene is one of the regulatory subunits. This subunit can be phosphorylated by the activated catalytic subunit. It may interact with various A-kinase anchoring proteins and determine the subcellular localization of cAMP-dependent protein kinase. This subunit has been shown to regulate protein transport from endosomes to the Golgi apparatus and further to the endoplasmic reticulum (ER). [provided by RefSeq, Jul 2008]

**GeneCards:** PRKAR2A (Protein Kinase CAMP-Dependent Type II Regulatory Subunit Alpha) is a Protein Coding gene. Diseases associated with PRKAR2A include Kallmann Syndrome. Among its related pathways are Beta-Adrenergic Signaling and Signaling by Hedgehog. Gene Ontology (GO) annotations related to this gene include *ubiquitin protein ligase binding* and *cAMP binding*.

*48. PSORS1C1*

**Entrez Gene Summary:** This gene is one of several genes thought to confer susceptibility to psoriasis and systemic sclerosis, located on chromosome 6 near the major histocompatibility complex (MHC) class I region. [provided by RefSeq, Sep 2011]

*49. RECQL4*

**Entrez Gene Summary:** The protein encoded by this gene is a DNA helicase that belongs to the RecQ helicase family. DNA helicases unwind double-stranded DNA into single-stranded DNAs and may modulate chromosome segregation. This gene is predominantly expressed in thymus and testis. Mutations in this gene are associated with Rothmund-Thomson, RAPADILINO and Baller-Gerold syndromes. [provided by RefSeq, Jan 2010]

**GeneCards:** RECQL4 (RecQ Like Helicase 4) is a Protein Coding gene. Diseases associated with RECQL4 include Baller-Gerold Syndrome and Rothmund-Thomson Syndrome. Among its related pathways are DNA Damage. Gene Ontology (GO) annotations related to this gene include *nucleic acid binding* and *annealing helicase activity*.

*50. REV3L*

**Entrez Gene Summary:** The protein encoded by this gene represents the catalytic subunit of DNA polymerase zeta, which functions in translesion DNA synthesis. The encoded protein can be found in mitochondria, where it protects DNA from damage. Defects in this gene are a cause of Mobius syndrome. [provided by RefSeq, Jan 2017]

**GeneCards:** REV3L (REV3 Like, DNA Directed Polymerase Zeta Catalytic Subunit) is a Protein Coding gene. Diseases associated with REV3L include Moebius Syndrome and Poland Syndrome. Among its related pathways are Translesion synthesis by Y family DNA polymerases bypasses lesions on DNA templateand Platinum drug resistance. Gene Ontology (GO) annotations related to this gene include *nucleic acid binding* and *4 iron, 4 sulfur cluster binding*.

*51. RPF2*

**GeneCards:** RPF2 (Ribosome Production Factor 2 Homolog) is a Protein Coding gene. Gene Ontology (GO) annotations related to this gene include rRNA binding.

*52. RPRML*

**GeneCards:** RPRML (Reprimo Like) is a Protein Coding gene. An important paralog of this gene is RPRM.

*53. SESN1*

**Entrez Gene Summary:** This gene encodes a member of the sestrin family. Sestrins are induced by the p53 tumor suppressor protein and play a role in the cellular response to DNA damage and oxidative stress. The encoded protein mediates p53 inhibition of cell growth by activating AMP-activated protein kinase, which results in the inhibition of the mammalian target of rapamycin protein. The encoded protein also plays a critical role in antioxidant defense by regenerating overoxidized peroxiredoxins, and the expression of this gene is a potential marker for exposure to radiation. Alternatively spliced transcript variants encoding multiple isoforms have been observed for this gene. [provided by RefSeq, Dec 2010]

**GeneCards:** SESN1 (Sestrin 1) is a Protein Coding gene. Diseases associated with SESN1 include Maxillary Cancer. Among its related pathways are Gene Expressionand DNA Damage Response. An important paralog of this gene is SESN3.

*54. SF3B1*

**Entrez Gene Summary:** This gene encodes subunit 1 of the splicing factor 3b protein complex. Splicing factor 3b, together with splicing factor 3a and a 12S RNA unit, forms the U2 small nuclear ribonucleoproteins complex (U2 snRNP). The splicing factor 3b/3a complex binds pre-mRNA upstream of the intron's branch site in a sequence independent manner and may anchor the U2 snRNP to the pre-mRNA. Splicing factor 3b is also a component of the minor U12-type spliceosome. The carboxy-terminal two-thirds of subunit 1 have 22 non-identical, tandem HEAT repeats that form rod-like, helical structures. Alternative splicing results in multiple transcript variants encoding different isoforms. [provided by RefSeq, Jul 2008]

**GeneCards:** SF3B1 (Splicing Factor 3b Subunit 1) is a Protein Coding gene. Diseases associated with SF3B1 include Myelodysplastic Syndrome and Acquired Idiopathic Sideroblastic Anemia. Among its related pathways are Activated PKN1 stimulates transcription of AR (androgen receptor) regulated genes KLK2 and KLK3and mRNA Splicing - Major Pathway. Gene Ontology (GO) annotations related to this gene include binding.

*55. SFXN2*

**GeneCards:** SFXN2 (Sideroflexin 2) is a Protein Coding gene. Gene Ontology (GO) annotations related to this gene include *cation transmembrane transporter activity* and *ion transmembrane transporter activity*. An important paralog of this gene is SFXN1.

*56. SLC25A20*

**Entrez Gene Summary:** This gene product is one of several closely related mitochondrial-membrane carrier proteins that shuttle substrates between cytosol and the intramitochondrial matrix space. This protein mediates the transport of acylcarnitines into mitochondrial matrix for their oxidation by the mitochondrial fatty acid-oxidation pathway. Mutations in this gene are associated with carnitine-acylcarnitine translocase deficiency, which can cause a variety of pathological conditions such as hypoglycemia, cardiac arrest, hepatomegaly, hepatic dysfunction and muscle weakness, and is usually lethal in new born and infants. [provided by RefSeq, Jul 2008]

**GeneCards:** SLC25A20 (Solute Carrier Family 25 Member 20) is a Protein Coding gene. Diseases associated with SLC25A20 include Carnitine-Acylcarnitine Translocase Deficiency and Hypoglycemia. Among its related pathways are Import of palmitoyl-CoA into the mitochondrial matrix and Regulation of lipid metabolism by Peroxisome proliferator-activated receptor alpha (PPARalpha).

*57. SMPD2*

**Entrez Gene Summary:** This gene encodes a protein which was initially identified as a sphingomyelinase based on sequence similarity between bacterial sphingomyelinases and a yeast protein. Subsequent studies showed that its biological function is less likely to be as a sphingomyelinase and instead as a lysophospholipase. [provided by RefSeq, Oct 2009]

**GeneCards:** SMPD2 (Sphingomyelin Phosphodiesterase 2) is a Protein Coding gene. Diseases associated with SMPD2 include Lipid Storage Disease and Coffin-Siris Syndrome 1. Among its related pathways are TNFR1 Pathway and p75(NTR)-mediated signaling. Gene Ontology (GO) annotations related to this gene include *sphingomyelin phosphodiesterase activity*.

*58. SOBP*

**Entrez Gene Summary:** The protein encoded by this gene is a nuclear zinc finger protein that is involved in development of the cochlea. Defects in this gene have also been linked to intellectual disability. [provided by RefSeq, Mar 2011]

**GeneCards:** SOBP (Sine Oculis Binding Protein Homolog) is a Protein Coding gene. Diseases associated with SOBP include Mental Retardation, Anterior Maxillary Protrusion, And Strabismus and Postcholecystectomy Syndrome. Gene Ontology (GO) annotations related to this gene include SUMO polymer binding. An important paralog of this gene is RAI2.

*59. SPPL2C*

**GeneCards:** SPPL2C (Signal Peptide Peptidase Like 2C) is a Protein Coding gene. Diseases associated with SPPL2C include Koolen-De Vries Syndrome. Gene Ontology (GO) annotations related to this gene include *protein homodimerization activity* and *aspartic-type endopeptidase activity*.

*60. STH*

**GeneCards:** STH (Saitohin) is a Protein Coding gene. Diseases associated with STH include Ascaridiasis and Trichuriasis.

*61. SYT1*

**Entrez Gene Summary:** The synaptotagmins are integral membrane proteins of synaptic vesicles thought to serve as Ca(2+) sensors in the process of vesicular trafficking and exocytosis. Calcium binding to synaptotagmin-1 participates in triggering neurotransmitter release at the synapse (Fernandez-Chacon et al., 2001 [PubMed 11242035]).[supplied by OMIM, Jul 2010]

**GeneCards:** SYT1 (Synaptotagmin 1) is a Protein Coding gene. Diseases associated with SYT1 include Mast-Cell Leukemia and Foodborne Botulism. Among its related pathways are Neurotransmitter Release Cycle and Transmission across Chemical Synapses. Gene Ontology (GO) annotations related to this gene include *calcium ion binding* and *transporter activity*.

*62. TBX21*

**Entrez Gene Summary:** This gene is a member of a phylogenetically conserved family of genes that share a common DNA-binding domain, the T-box. T-box genes encode transcription factors involved in the regulation of developmental processes. This gene is the human ortholog of mouse Tbx21/Tbet gene. Studies in mouse show that Tbx21 protein is a Th1 cell-specific transcription factor that controls the expression of the hallmark Th1 cytokine, interferon-gamma (IFNG). Expression of the human ortholog also correlates with IFNG expression in Th1 and natural killer cells, suggesting a role for this gene in initiating Th1 lineage development from naive Th precursor cells. [provided by RefSeq, Jul 2008]

**GeneCards:** TBX21 (T-Box 21) is a Protein Coding gene. Diseases associated with TBX21 include Asthma, Nasal Polyps, And Aspirin Intolerance and Genital Herpes. Among its related pathways are Development and heterogeneity of the ILC family and Glucocorticoid receptor regulatory network. Gene Ontology (GO) annotations related to this gene include *DNA-binding transcription factor activity* and *transcription regulatory region DNA binding*.

*63. TFDP2*

**Entrez Gene Summary:** The gene is a member of the transcription factor DP family. The encoded protein forms heterodimers with the E2F transcription factors resulting in transcriptional activation of cell cycle regulated genes. Alternative splicing results in multiple transcript variants. [provided by RefSeq, May 2010]

**GeneCards:** TFDP2 (Transcription Factor Dp-2) is a Protein Coding gene. Among its related pathways are TP53 Regulates Transcription of Cell Cycle Genes and Transcriptional activity of SMAD2/SMAD3-SMAD4 heterotrimer. Gene Ontology (GO) annotations related to this gene include *DNA-binding transcription factor activity* and *protein domain specific binding*.

*64. TMEM180*

**GeneCards:** MFSD13A (Major Facilitator Superfamily Domain Containing 13A) is a Protein Coding gene.

*65. USP19*

**Entrez Gene Summary:** Protein ubiquitination controls many intracellular processes, including cell cycle progression, transcriptional activation, and signal transduction. This dynamic process, involving ubiquitin conjugating enzymes and deubiquitinating enzymes, adds and removes ubiquitin. Deubiquitinating enzymes are cysteine proteases that specifically cleave ubiquitin from ubiquitin-conjugated protein substrates. This protein is a ubiquitin protein ligase and plays a role in muscle wasting. Alternatively spliced transcript variants encoding different isoforms have been found for this gene. [provided by RefSeq, May 2017]

**GeneCards:** USP19 (Ubiquitin Specific Peptidase 19) is a Protein Coding gene. Diseases associated with USP19 include Nephrotic Syndrome, Type 5, With Or Without Ocular Abnormalities and Pierson Syndrome. Among its related pathways are Metabolism of proteins and Ubiquitin-Proteasome Dependent Proteolysis. Gene Ontology (GO) annotations related to this gene include *cysteine-type endopeptidase activity* and *thiol-dependent ubiquitin-specific protease activity*.

*66. VARS2*

**Entrez Gene Summary:** This gene encodes a mitochondrial aminoacyl-tRNA synthetase, which catalyzes the attachment of valine to tRNA(Val) for mitochondrial translation. Mutations in this gene cause combined oxidative phosphorylation deficiency-20, and are also associated with early-onset mitochondrial encephalopathies. Alternative splicing of this gene results in multiple transcript variants. [provided by RefSeq, Aug 2014]

**GeneCards:** VARS2 (Valyl-TRNA Synthetase 2, Mitochondrial) is a Protein Coding gene. Diseases associated with VARS2 include Combined Oxidative Phosphorylation Deficiency 20 and Combined Oxidative Phosphorylation Deficiency. Among its related pathways are tRNA Aminoacylation and Gene Expression. Gene Ontology (GO) annotations related to this gene include nucleotide binding and aminoacyl-tRNA editing activity.

*67. WNT3*

**Entrez Gene Summary:** The WNT gene family consists of structurally related genes which encode secreted signaling proteins. These proteins have been implicated in oncogenesis and in several developmental processes, including regulation of cell fate and patterning during embryogenesis. This gene is a member of the WNT gene family. It encodes a protein which shows 98% amino acid identity to mouse Wnt3 protein, and 84% to human WNT3A protein, another WNT gene product. The mouse studies show the requirement of Wnt3 in primary axis formation in the mouse. Studies of the gene expression suggest that this gene may play a key role in some cases of human breast, rectal, lung, and gastric cancer through activation of the WNT-beta-catenin-TCF signaling pathway. This gene is clustered with WNT15, another family member, in the chromosome 17q21 region. [provided by RefSeq, Jul 2008]

**GeneCards:** WNT3 (Wnt Family Member 3) is a Protein Coding gene. Diseases associated with WNT3 include Tetraamelia Syndrome 1 and Tetra-Amelia Syndrome. Among its related pathways are Hippo signaling pathway and Nanog in Mammalian ESC Pluripotency. Gene Ontology (GO) annotations related to this gene include signaling receptor binding and frizzled binding.

### SUPPLEMENTARY FIGURES

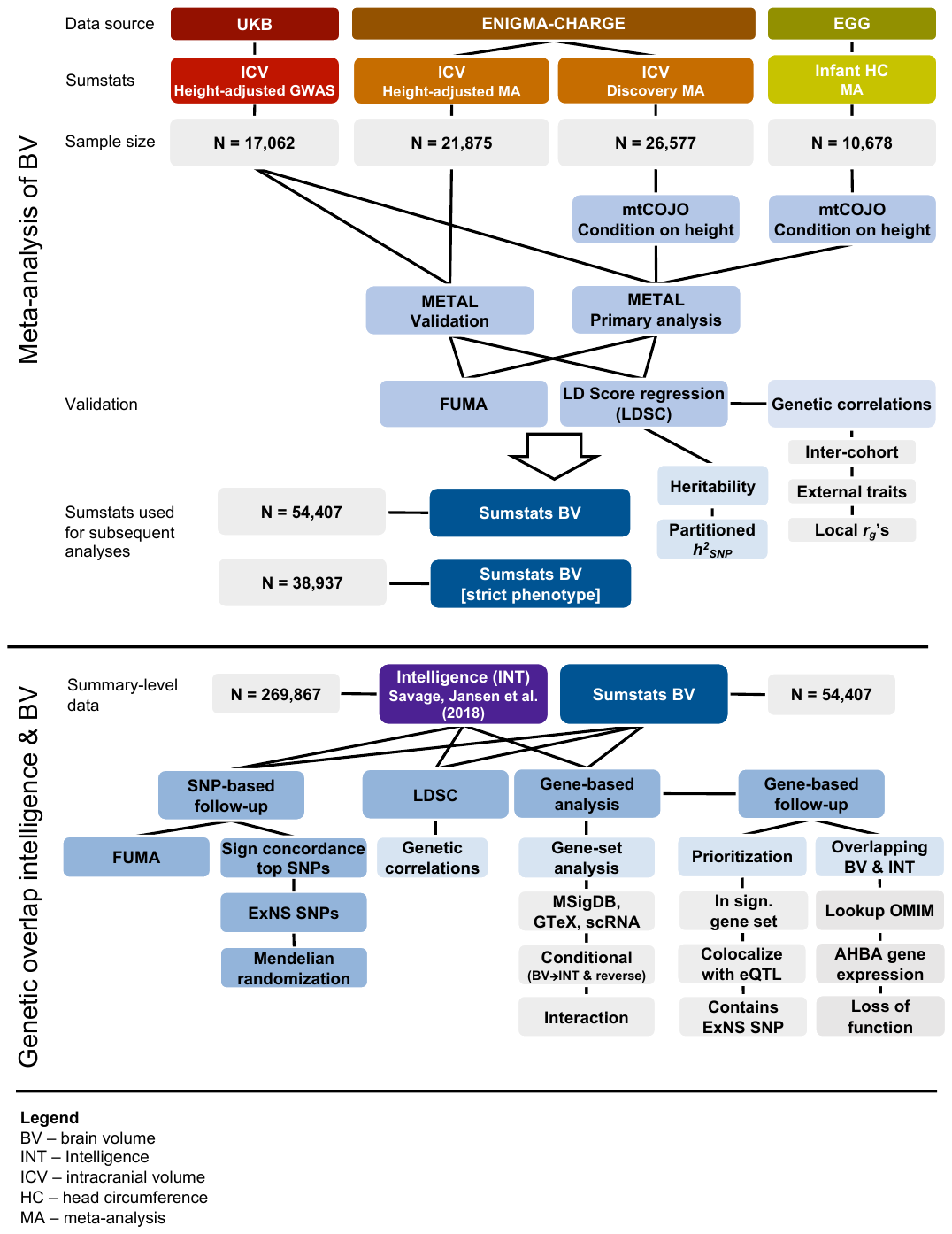

**Supplementary Fig. 1. Flowchart of the GWAS meta-analyses of brain volume.** Schematic representation of the analyses conducted in the current study. ICV = intracranial volume; HC = head circumference; BV = brain volume; INT = Intelligence.

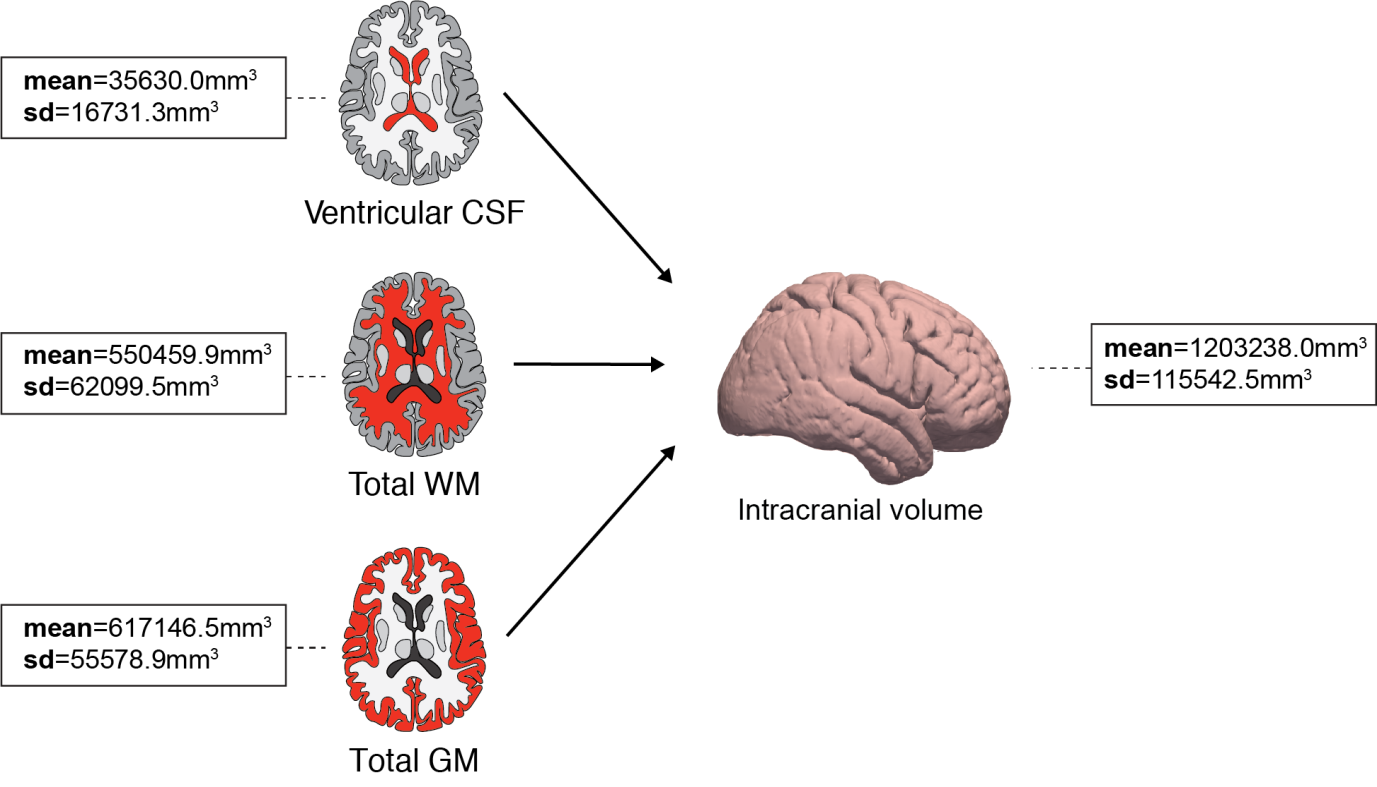
­­

**Supplementary Fig. 2**. **Distribution of brain structures included in the brain volume phenotype in 17,062 individuals of the UK Biobank.** Mean and standard deviation (sd) of the three brain phenotypes that were combined in the brain volume measure. Brain volume means and sd’s are expressed in cubic millimeters (mm^3^). CSF=cerebrospinal fluid; WM=white matter; GM=gray matter.

**Supplementary Fig. 3. Manhattan and QQ-plots of the individual genome-wide association analyses of brain volume.** Manhattan plot showing negative log_10_-transformed *P-*values on the *y*-axis and genomic position on the *x*-axis, and QQ-plots showing the observed versus the expected *P*-value distribution. Results are shown for **(a)** the GWAS of brain volume in UK Biobank (N=17,062 individuals), **(b)** the GWAS of intracranial volume in the ENIGMA-CHARGE consortium (N=26,577 individuals), and **(c)** the GWAS of head circumference in the EGG Consortium (N=10,768 individuals). The data for the latter two samples was corrected for height using mtCOJO (see **Online Methods**).

**Supplementary Fig. 4. Regional association plots of the meta-analyzed SNP results for BV.** Regional plots were created using LocusZoom for all genomic loci identified in the GWAS meta-analysis of BV (N=54,407 individuals). The *y*-axis represents the negative log_10_-transformed SNP *P*-values and the *x*-axis indicates the genomic position. Level of LD between a SNP and the lead SNP (purple) is indicated by the color.
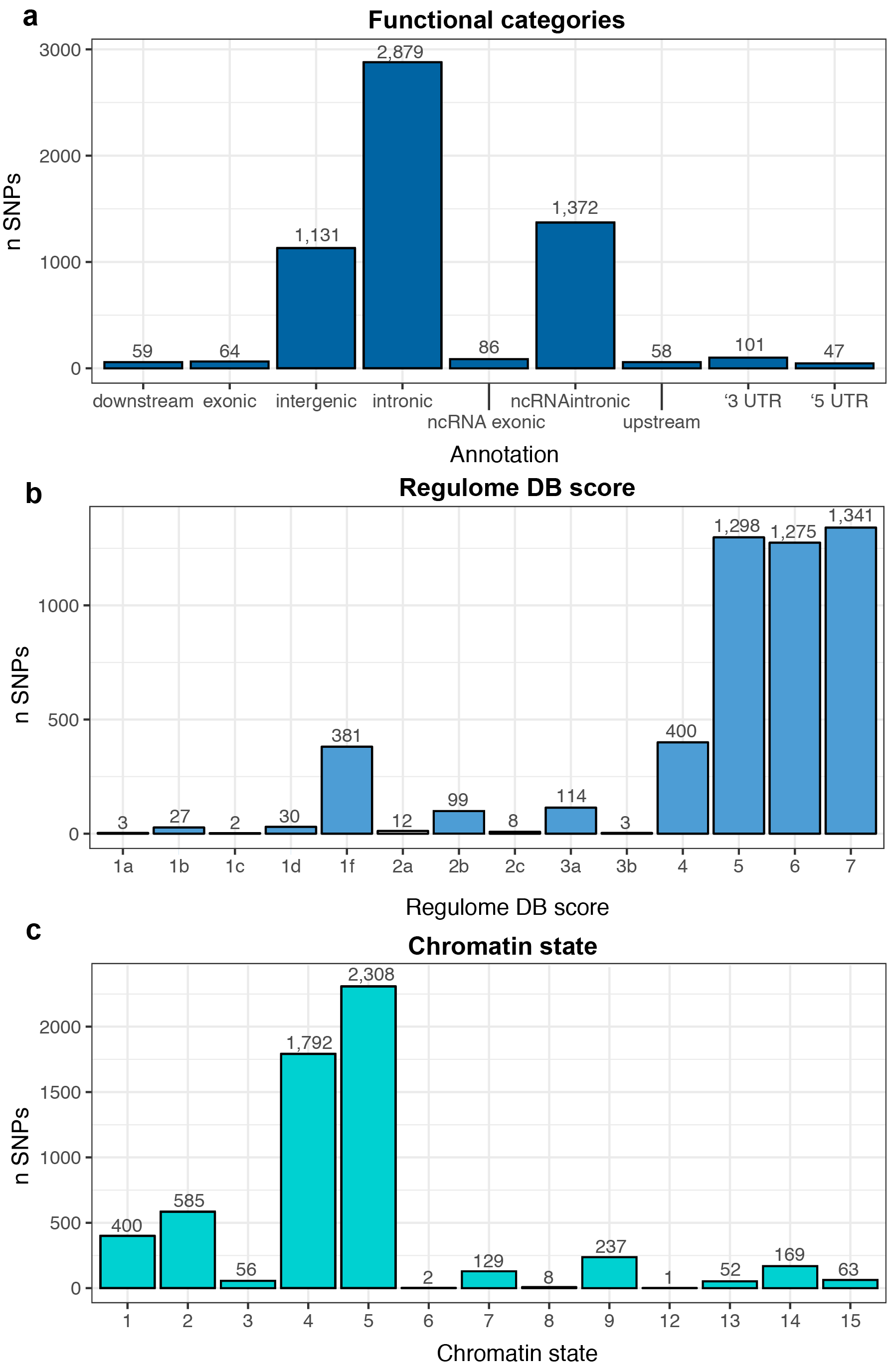

**Supplementary Fig. 5. Functional annotation of the GWAS meta-analysis results of brain volume in 54,407 individuals.** Functional annotation was carried out in FUMA on all 5,802 *candidate* SNPs (LD > 0.6 with an independent significant SNP and *P-*value of < 1×10^-5^). **(a)** Number of SNPs annotated to the functional SNP categories. **(b)** Number of SNPs in each Regulome DB score category. A lower score represents a higher likelihood that the SNP has a regulatory function. (**c**) Number of SNPs in each of 15 possible chromatin states.

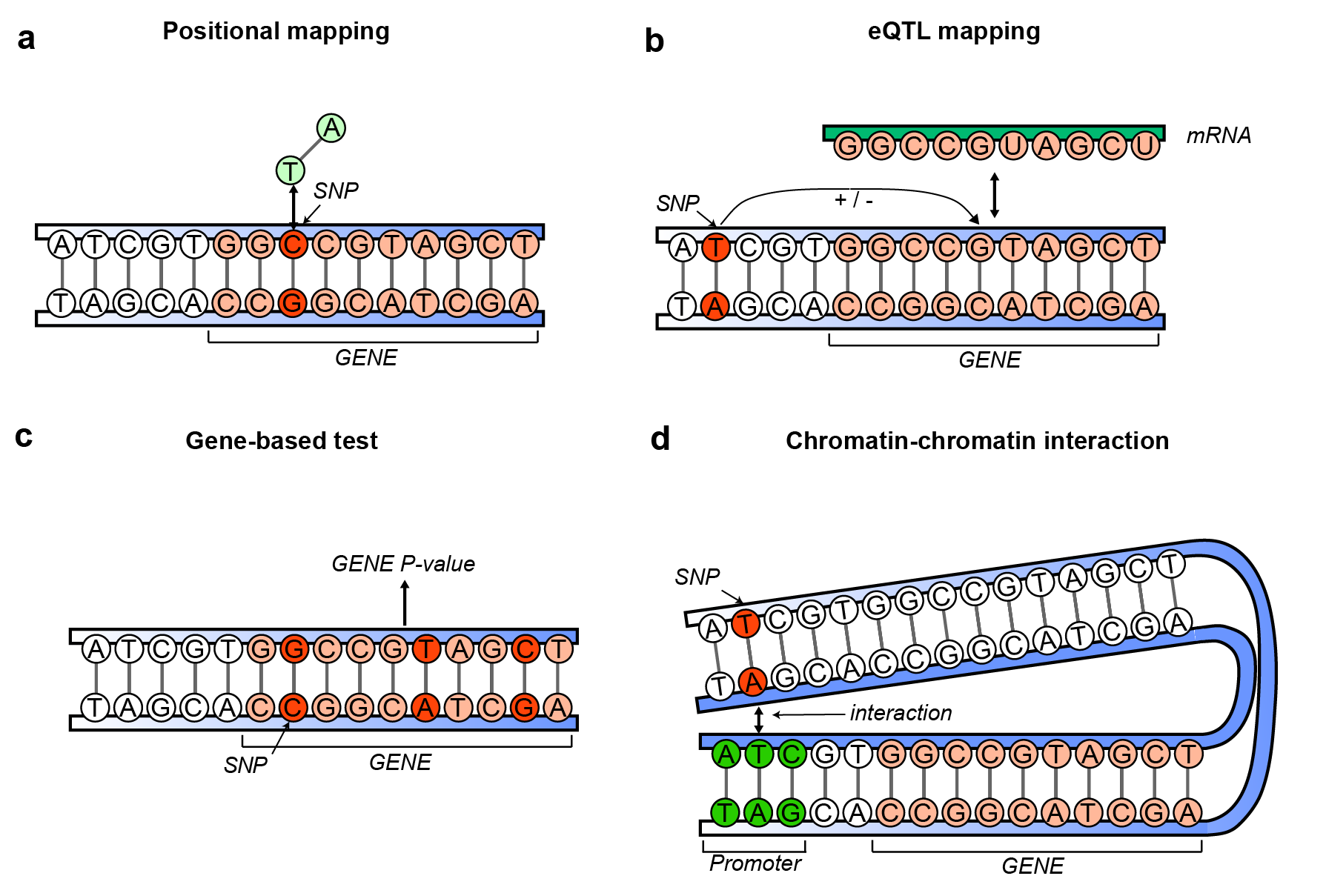

**Supplementary Fig. 6. Gene-mapping methods that were used to map SNPs from GWAS to genes.** Schematic representation of gene-mapping efforts used in the GWAS follow-up analyses: **(a)** positional gene-mapping based position of genome-wide significant (GWS) SNPs within the physical boundaries (or within proximity, i.e., within 10 kb window) of a gene, **(b)** gene-mapping through eQTL, where a GWS SNP is known to influence the expression levels of a gene, **(c)** gene-based association testing (MAGMA), where SNP association *P-*values within a gene are combined into a gene-based *P-*value, and **(d)** gene-mapping through chromatin-chromatin interaction, where GWS SNPs physically interact with genes through the 3D structure of the genome.

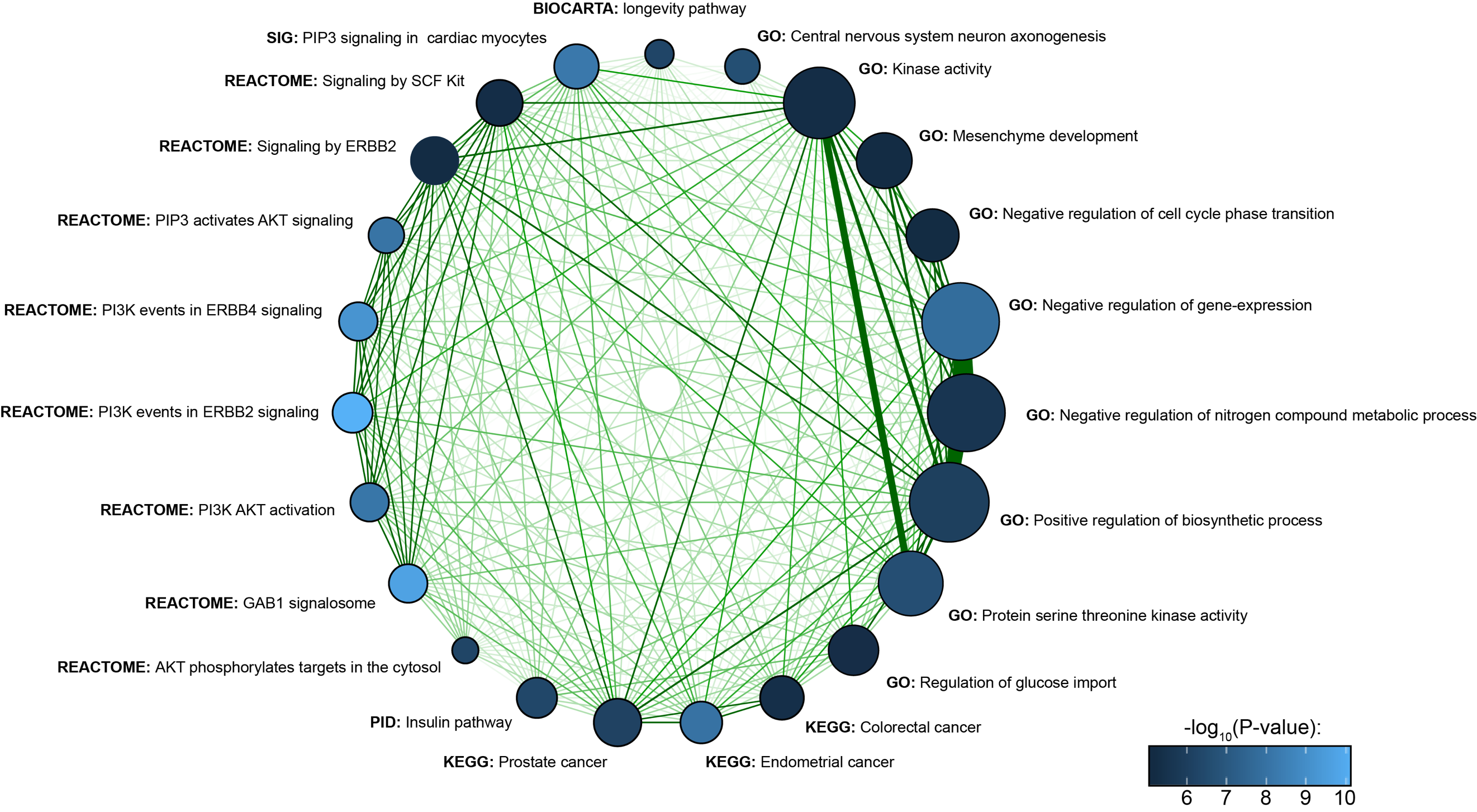

**Supplementary Fig. 7. Significant gene sets in the GWAS meta-analysis of brain volume.** Gene-set analysis was performed in MAGMA using gene-based *P-*values as input. Gene sets are shown that passed the stringent correction for multiple testing (*P* < 0.05/7,864 = 6.36×10^-6^). The number of genes in the gene set are represented by the size of the circle, the -log_10_-transformed *P-*value of the gene set by the color of the circle, and the number of genes that overlap between gene sets by the thickness of the connections.

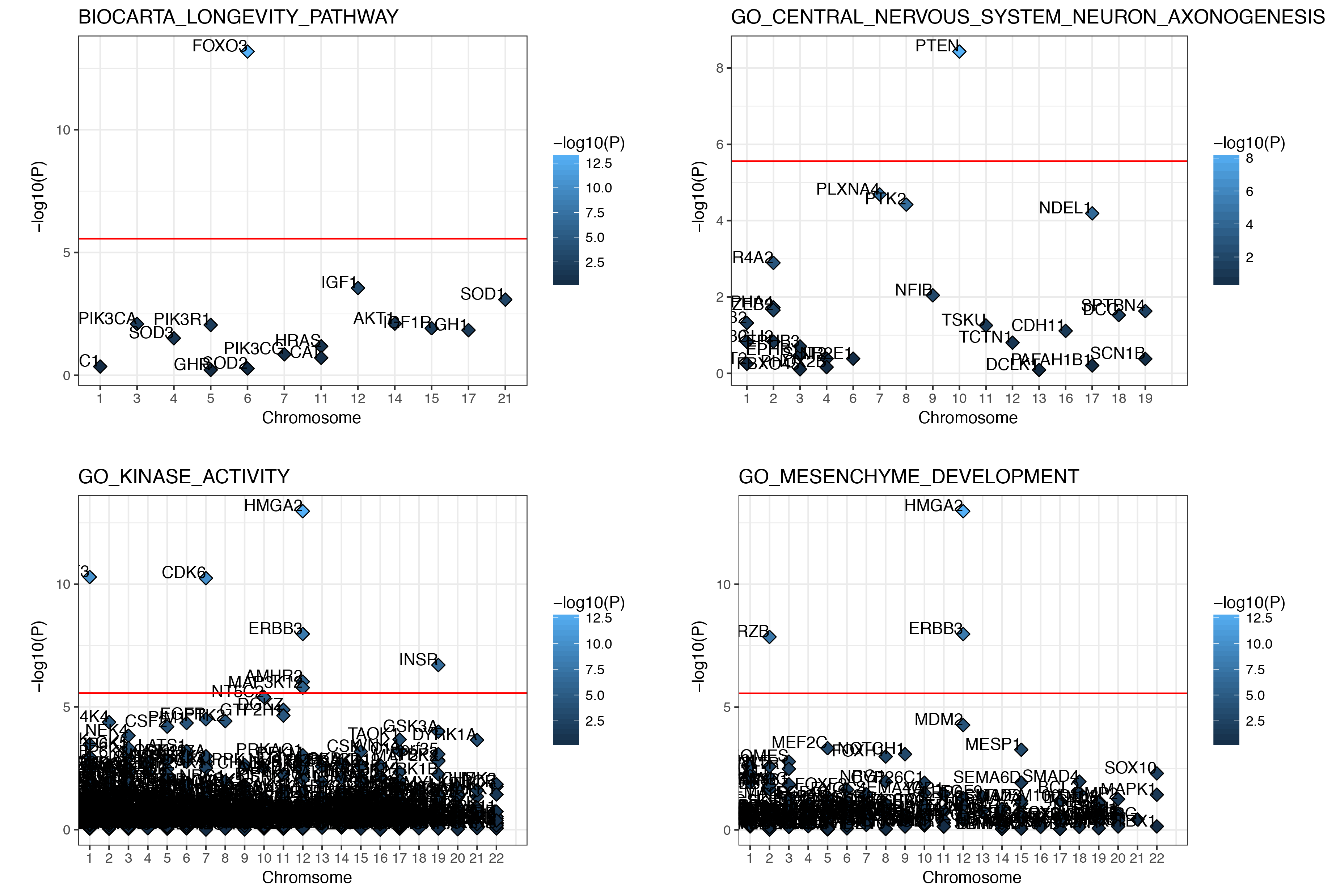

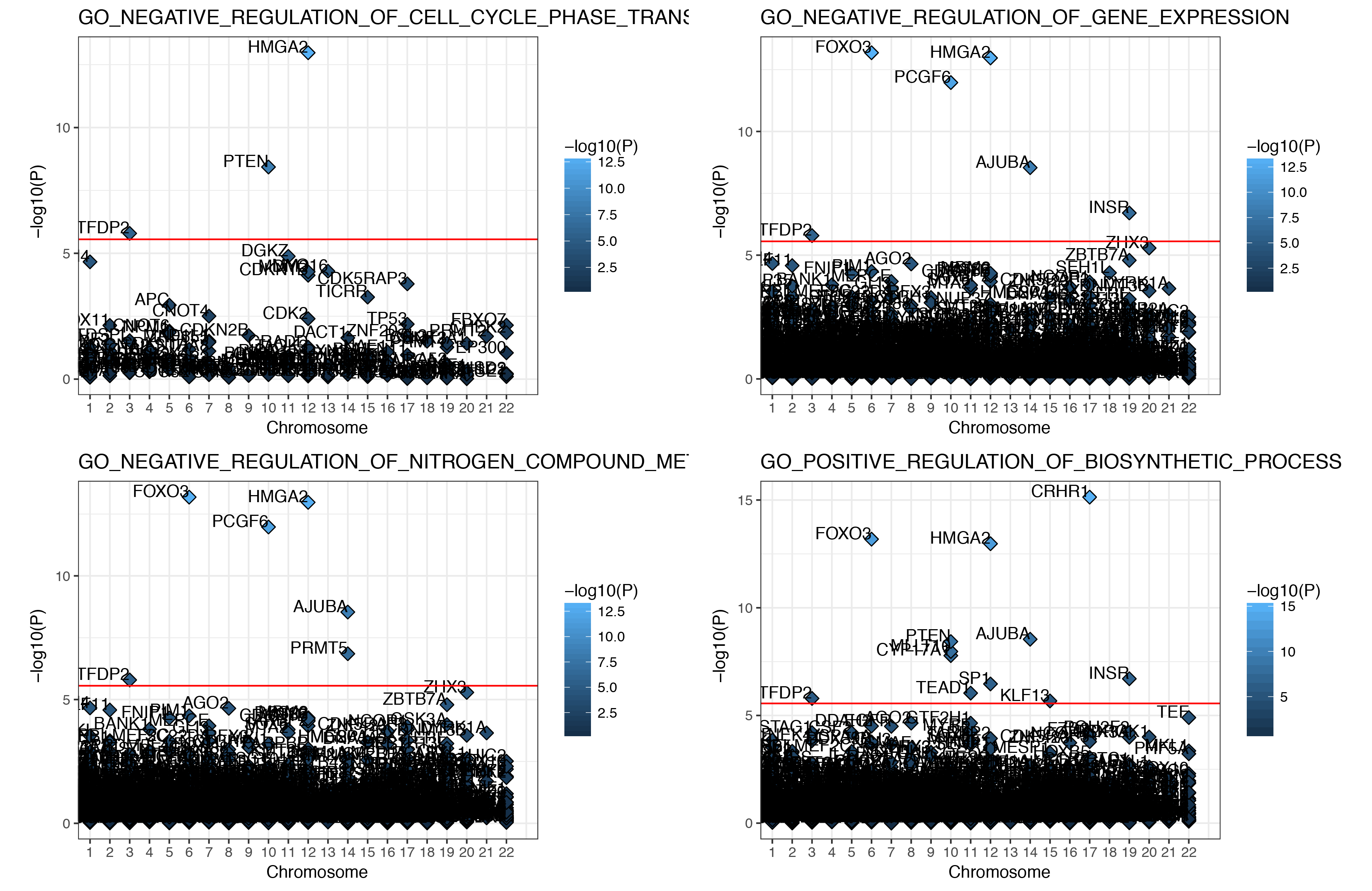

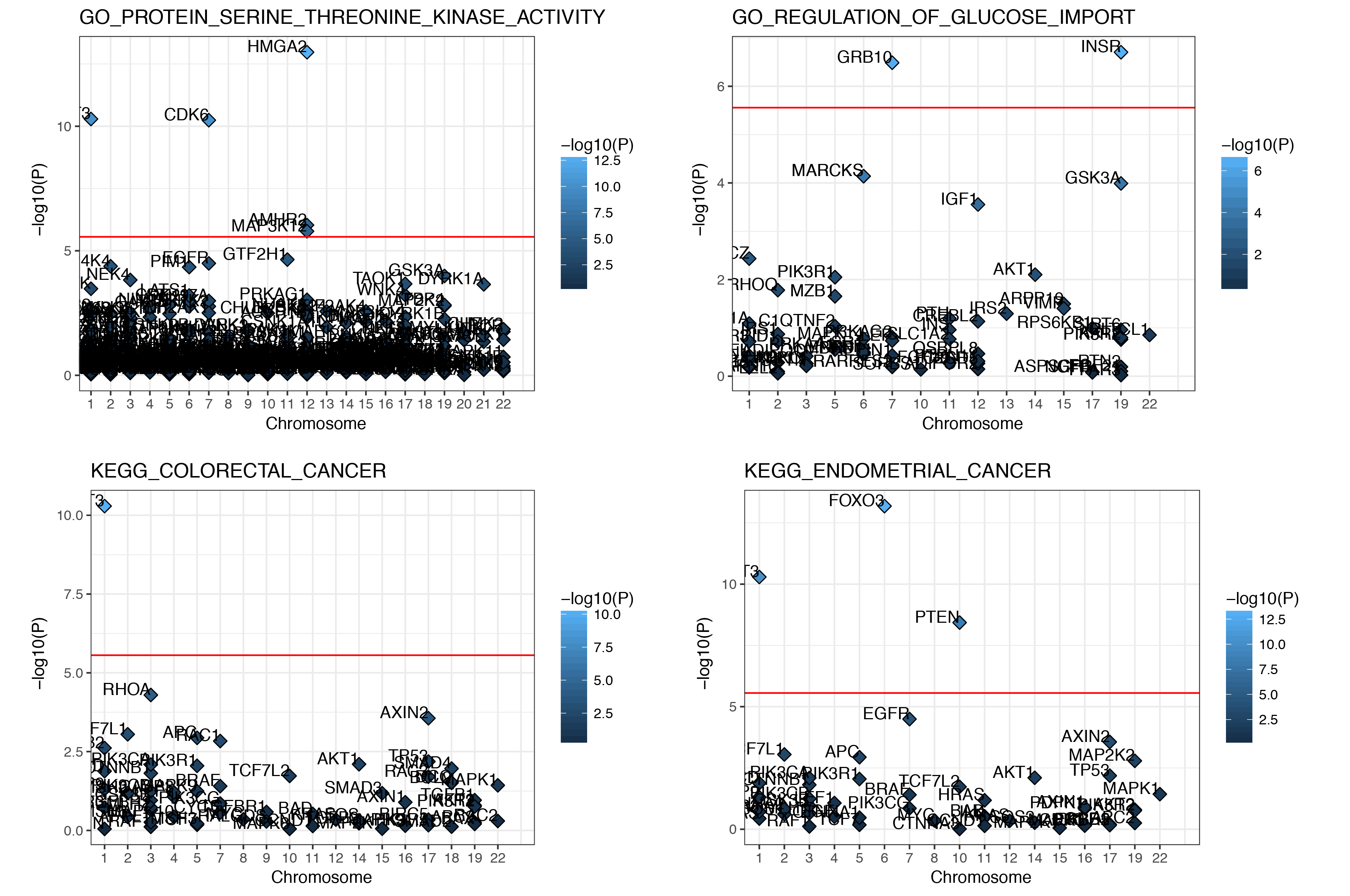

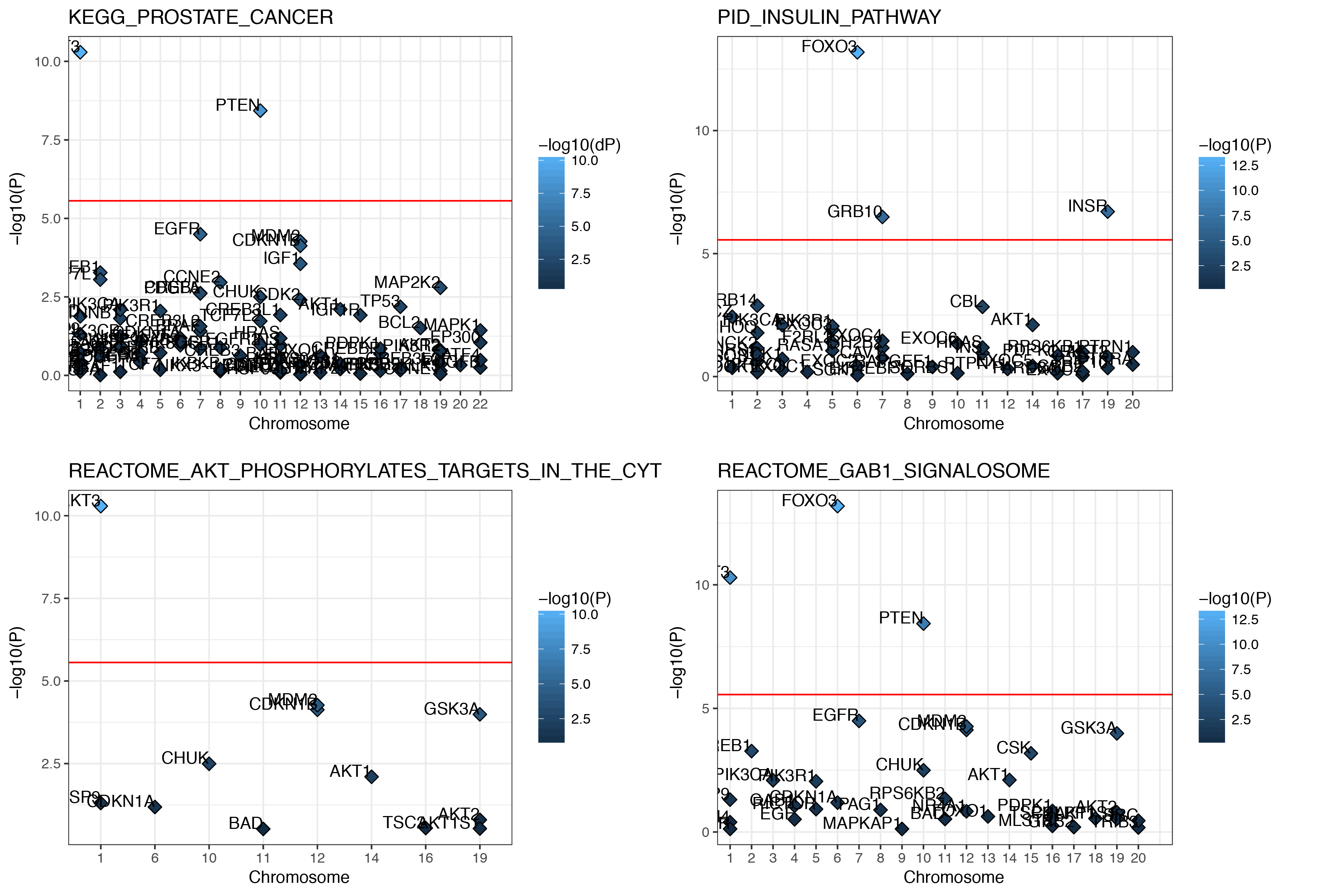

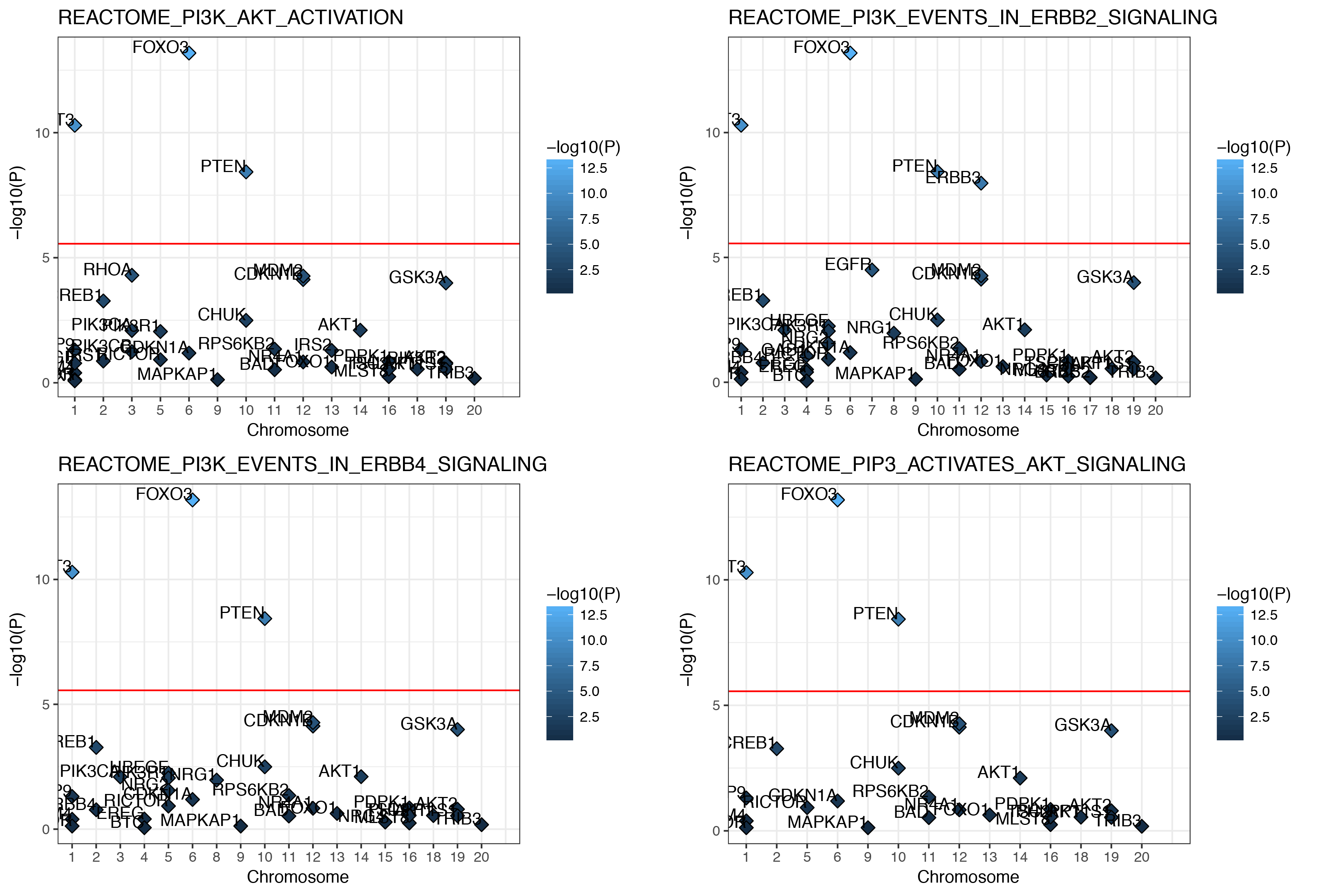

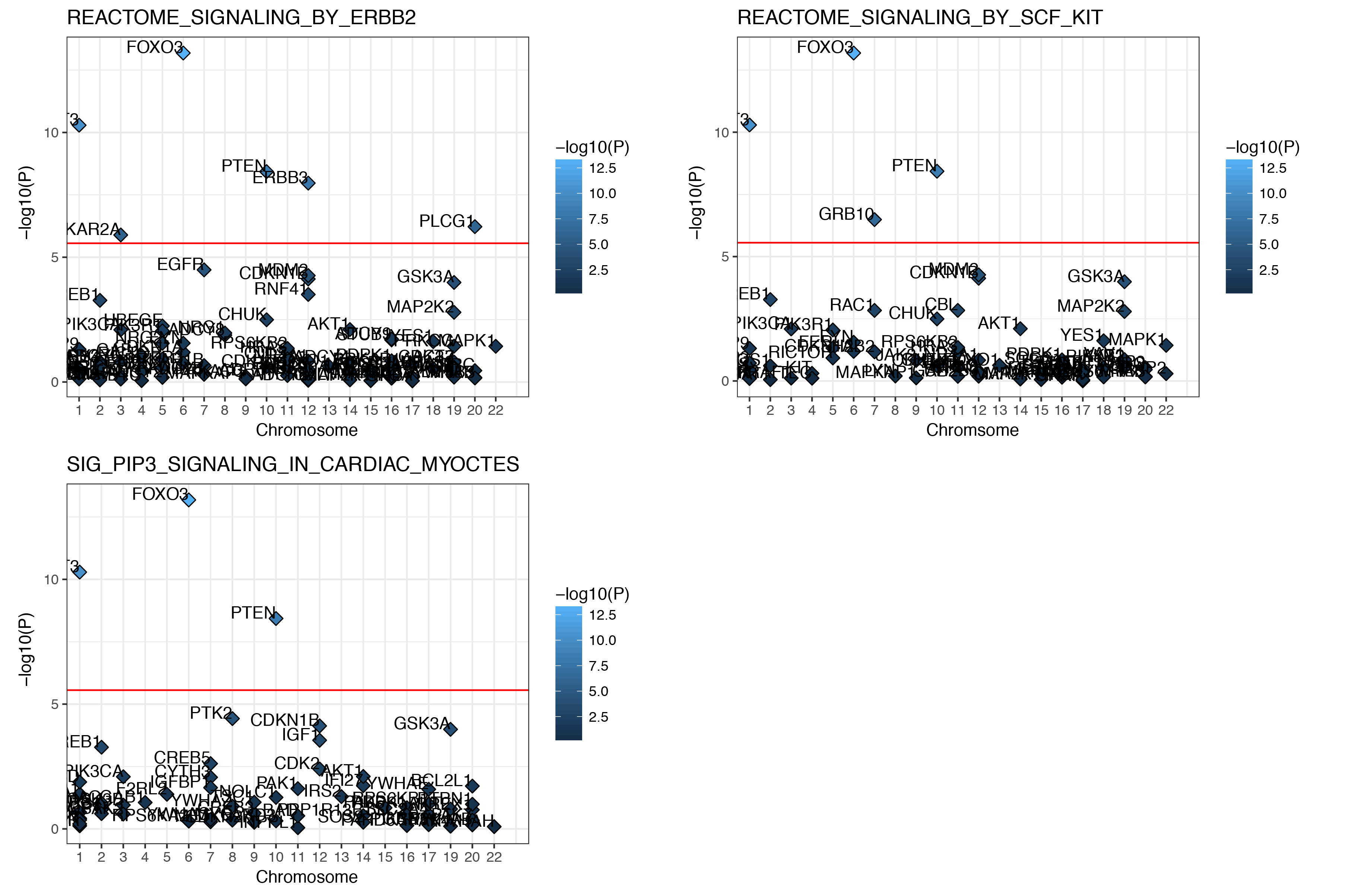
**Supplementary Fig. 8. Gene associations within each of the significant gene set.** Gene-based *P-*values are shown for all genes that are part of one of the 23 gene sets significantly associated with BV in MAGMA (gene-sets *P* < 0.05/7,864 = 6.36×10^-6^). The chromosomal location of each gene within the gene set is shown on the x-axis, whereas the -log_10_-transformed *P-*value of the gene in the gene-based test in MAGMA is shown on the y-axis. Genes are annotated by gene name.

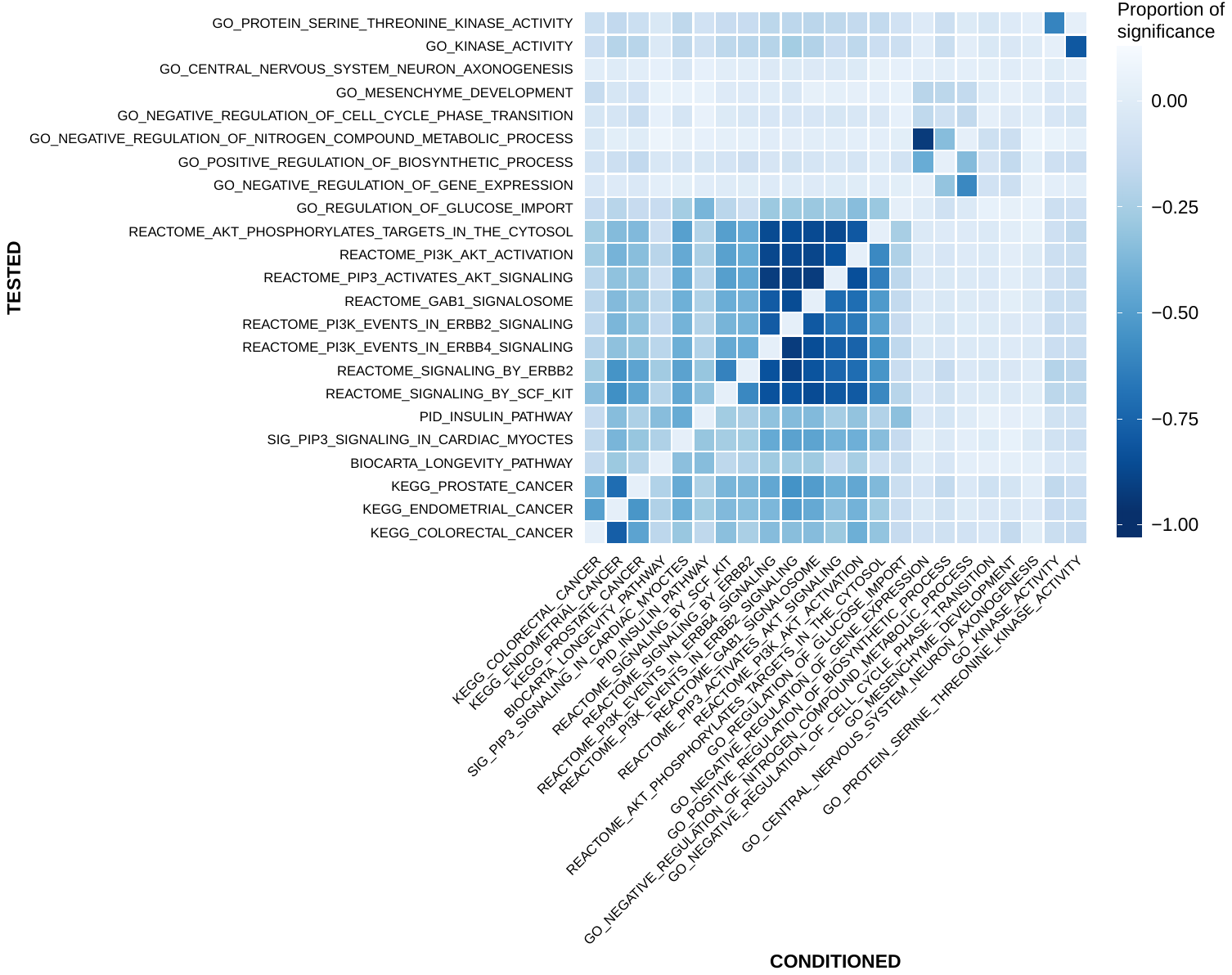

**Supplementary Fig. 9. Conditional gene-set analysis**. Heatmap of the pairwise conditional gene-set analyses of the 23 gene-sets significant for BV. Off-diagonal entries represents the scaled difference between negative log_10_-transformed *P-*values of conditional minus marginal gene-set association *P*-values (Proportion of significance = (-log10(P_cond) minus -log10(P_marg)) / -log10(P_marg); with more negative values indicating that the marginal P-value is lower than the conditional *P*-value. Gene-sets on the y-axis were conditioned on the gene-set on the x-axis. All gene-set analyses were performed in MAGMA.

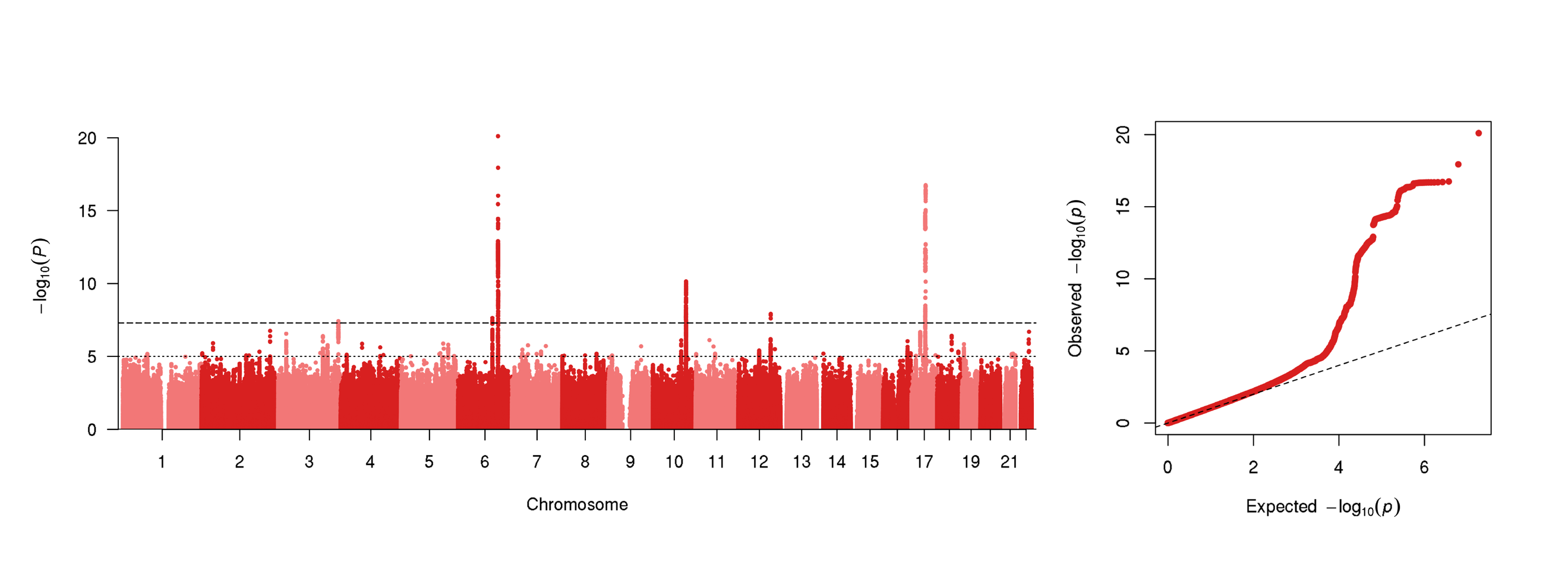

**Supplementary Fig. 10. Manhattan and QQ-plot of ICV in the subset of ENIGMA-CHARGE data (N = 21,875 individuals).** Manhattan plot showing negative log_10_-transformed *P-*values on the *y*-axis and genomic position on the *x*-axis, and QQ-plots showing the observed versus the expected *P*-value distribution. This figure shows data for the subset for which height was included as a covariate in the original analysis by the ENIGMA-CHARGE collaboration (rather than corrected for height post-hoc using mtCOJO).

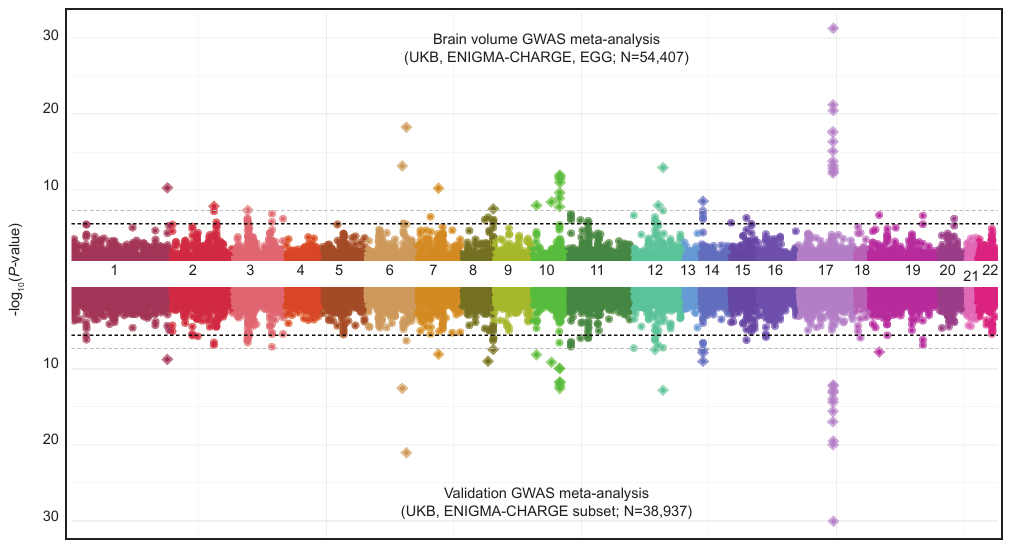

**Supplementary Fig. 11. Miami-plot of the gene-based association test in the main GWAS meta-analysis of BV and the validation GWAS meta-analysis.** The *y*-axis of the Miami plot shows the negative log_10_-transformed SNP *P-*value of each gene, and the *x*-axis indicates the genomic position. The top part is based on the main GWAS meta-analysis of brain volume in UK Biobank, ENIGMA-CHARGE and EGG (N=54,407), where the GWAS summary statistics of EGG and ENIGMA-CHARGE were corrected for height using mtCOJO. The bottom part shows the validation meta-analysis that strictly included ICV data from UK Biobank and ENIGMA-CHARGE (N=38,937). The dashed black line indicates the genome-wide significance threshold (*P* < 2.75×10^-6^). Genes indicated with a diamond are considered GWS.

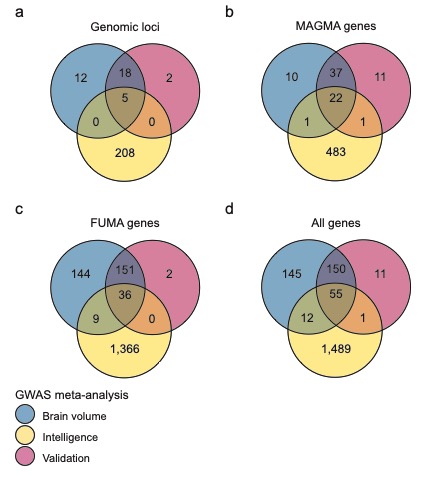

**Supplementary Fig. 12. Venn diagram showing overlap in number of loci and genes between the three meta-analyses.** Overlap in results between the main brain volume (BV) GWAS meta-analysis (blue circle), intelligence (yellow circle) and the validation ICV GWAS meta-analysis (red circle) is shown for **(a**) physically overlapping loci, (**b)** overlap in significant genes in MAGMA, **(c)** overlap in genes mapped by FUMA, **(d)** overlap in all unique genes mapped by either method.

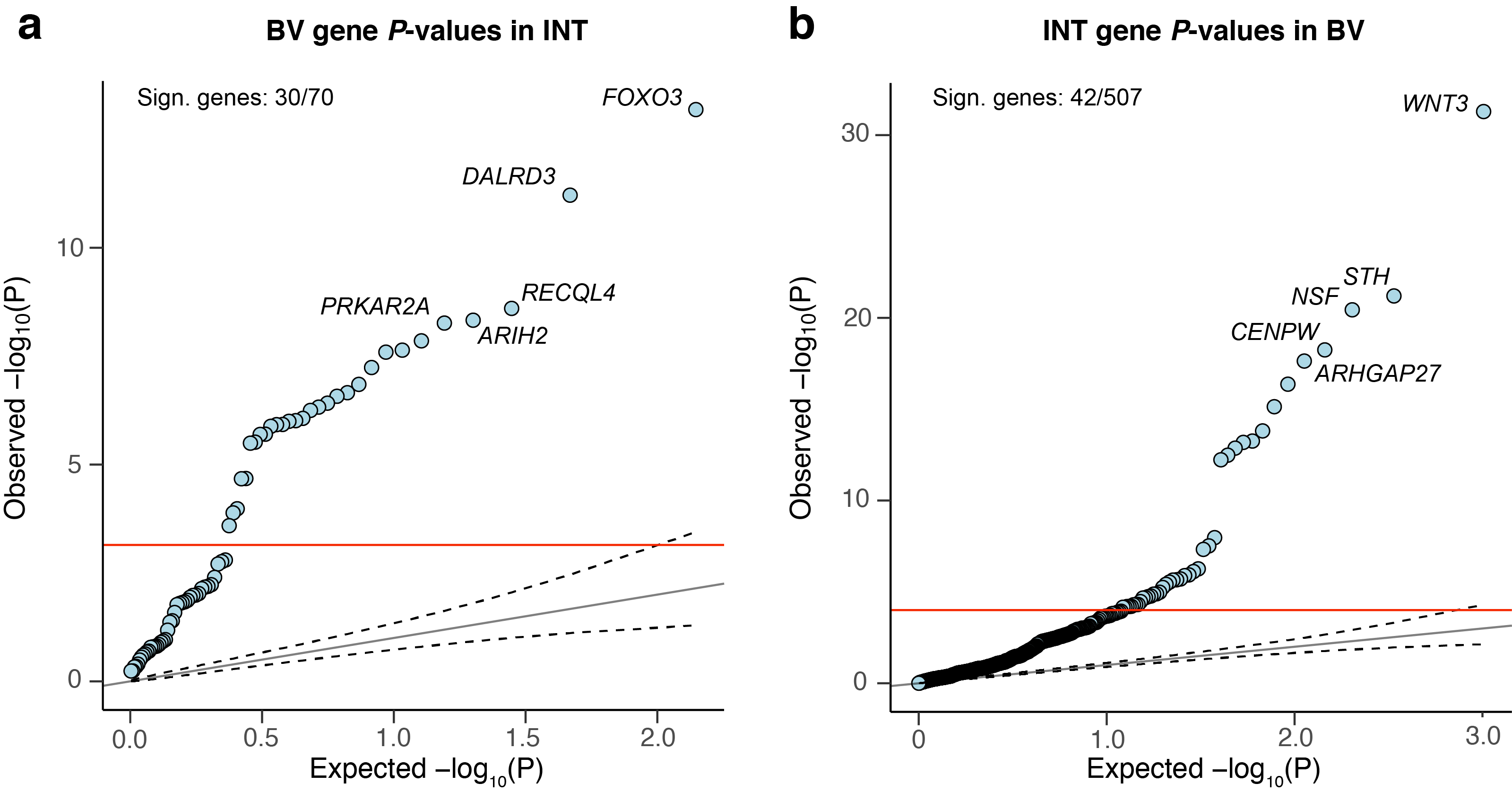

**Supplementary Fig. 13. QQ-plots of gene *P-*values of significant BV genes in intelligence and vice versa.** The *x*-axis shows the expected *P-*value distribution and the *y*-axis the observed cross-trait *P-*value of the gene. Results are shown for **(a)** *P-*values of significant genes in the brain volume (BV) meta-analysis in the gene-based test based on the meta-analysis of intelligence (INT), and **(b)** *P-*values of significant genes in the meta-analysis of intelligence in the gene-based test based on the BV GWAS meta-analysis. The dashed lines represent the upper and lower limit of the 95% confidence interval around the observed-*P-*value-equals-expected-*P-*value line. The red line indicates the Bonferroni-corrected threshold of the number of replicated genes in each gene-based association test (**a**: *P*<0.05/70 genes = 7.14×10^-4^; **b**: *P*<0.05/507=9.86×10^-5^). The number in the upper-left corner indicates the number of genes that were significant in BV and were replicated in INT, and vice versa.
